# The metabolic enzyme GYS1 condenses with NONO/p54^nrb^ in the nucleus to spatiotemporally regulate glycogenesis and myogenic differentiation

**DOI:** 10.1101/2025.01.19.633807

**Authors:** Shujun Peng, Canrong Li, Yifan Wang, Yuguo Yi, Xinyu Chen, Yujing Yin, Fan Yang, Fengzhi Chen, Yingyi Ouyang, Haolun Xu, Baicheng Chen, Haowen Shi, Qingrun Li, Yu Zhao, Lin Feng, Zhenji Gan, Xiaoduo Xie

## Abstract

Accumulating evidence indicates that metabolic enzymes can directly couple metabolic signals to transcriptional adaptation and cell differentiation. Glycogen synthase 1 (GYS1), the key metabolic enzyme for glycogenesis, is a nucleocytoplasmic shuttling protein compartmentalized in the cytosol and nucleus. However, the spatiotemporal regulation and biological function of nuclear GYS1 (nGYS1) microcompartments remain unclear. Here, we show that GYS1 dynamically reorganizes into nuclear condensates under conditions of glycogen depletion or transcription inhibition. nGYS1 complexes with the transcription factor NONO/p54^nrb^ and undergoes liquid-liquid phase separation to form biomolecular condensates, leading to its nuclear retention and inhibition of glycogen biosynthesis. Compared to their wild-type littermates, Nono-deficient mice exhibit exercise intolerance, higher muscle glycogen content, and smaller myofibers. Additionally, Gys1 or Nono deficiency prevents C2C12 differentiation and cardiotoxin-induced muscle regeneration in mice. Mechanistically, nGYS1 and NONO co-condense with the myogenic transcription factor MyoD and preinitiation complex (PIC) proteins to form transcriptional condensates, driving myogenic gene expression during myoblast differentiation. These results reveal the spatiotemporal regulation and subcellular function of nuclear GYS1 condensates in glycogenesis and myogenesis, providing mechanistic insights into glycogenoses and muscular dystrophy.

## Introduction

Glycogen synthase 1 (GYS1) is the rate-limiting enzyme for glycogen biosynthesis in various tissues, including muscle and the brain [1, 2]. Dysregulation of GYS1 leads to systemic metabolic disorders, such as glycogen storage diseases (GSDs or glycogenoses) [3, 4]. GYS1 assembles with glycogen into a supramolecular glycogen-protein complex known as a glycogen granule or glycosome. Glycogen metabolic proteins, including GYS1 and the initiator enzyme glycogenin 1 (GYG1), have been identified as components of the glycosome [5, 6]. GYS1 is a nucleocytoplasmic shuttling protein, compartmentalized in the cytosol and nucleus [7, 8]. This tetrameric enzyme is regulated by glucose-6-phosphate (G6P) allosteric binding and kinase-mediated multisite phosphorylation in response to metabolic signals [1, 2]. GYS1 belongs to the glycosyltransferase family B (GT-B) and contains distinct N- and C-terminal Rossmann-fold (Rf) domains [9, 10], which confer multiple functions on metabolic enzymes, including nucleotide sugar donor binding, RNA binding, and ribonuclease activity [11, 12]. Although GYS1 has been reported to associate with rRNA and mRNA [13, 14], the specific RNA species it binds and the biological implications remain unclear.

The formation of intracellular protein bodies for metabolic enzymes, known as membrane-less organelles (MLOs), is an evolutionarily conserved biological process prevalent in prokaryotes and eukaryotes. These MLOs typically self-assemble as protein fibers or foci under specific metabolic stresses [15, 16]. The formation of MLOs is primarily driven by enzymes with intrinsically disordered regions (IDRs) and DNA- or RNA-binding domains through protein-phase separation (PPS), leading to the dynamic formation of intracellular bodies that exhibit solid, gel, or liquid-like properties [17, 18]. MLOs allow for efficient control of metabolic pathways, facilitate substrate channeling, and minimize side reactions. Moreover, they enable spatial and temporal regulation of enzymatic activities, contributing to metabolic homeostasis in response to specific stimuli or stresses [15]. Previous studies have identified numerous phase-separated MLOs, including stress granules, nucleoli, processing bodies, purinosomes, G bodies, and nuclear bodies (NBs) [19–22]. The glycosome has long been recognized as an MLO in the cytosol [5, 6], possibly through a phase separation mechanism [23]. However, whether the GYS1 protein also participates in the formation of MLOs and how subcellular GYS1 protein bodies are spatiotemporally regulated remain to be explored.

The non-POU domain-containing octamer-binding protein NONO (also known as p54^nrb^) is an X-linked gene product belonging to the highly conserved Drosophila behavior/human splicing (DBHS) protein family [24, 25]. NONO is a multifunctional scaffold protein with DNA- and RNA-binding domains, exhibiting the property of protein liquid-liquid phase separation (LLPS) [26–28]. It functions as a transcription factor (TF) or RNA-binding protein (RBP), regulating gene transcription and post-transcriptional RNA processing [25, 29]. DBHS proteins, along with other RBPs such as fused in sarcoma (FUS), undergo phase separation with the long non-coding RNA (lncRNA) nuclear paraspeckle assembly transcript 1 (NEAT1) isoform 2 (NEAT1_2) to form stress-sensing nuclear bodies known as paraspeckles [30, 31]. NONO and other DBHS proteins also have non-paraspeckle functions as multifaceted regulators in DNA damage repair, metabolism, transcription, and cell differentiation [25, 29, 32].

Skeletal muscle possesses remarkable myogenic and regenerative capabilities conferred by muscle stem cells (MuSCs) or satellite cells in response to injury or metabolic stresses [33, 34]. Myoblast differentiation is crucial for muscle regeneration and is primarily governed by the activation of myogenic differentiation 1 (MyoD) and myogenin (Myog), followed by the expression of myosin heavy chain (MyHC) genes [35, 36]. MyoD is the master TF that regulates myogenesis [36, 37]. Recent studies suggest that TFs, coactivators, and RNA polymerase II (Pol II) condense into transcriptional condensates at genomic loci through IDR-mediated liquid–liquid phase separation (LLPS). These condensates effectively activate gene transcription, particularly during progenitor cell differentiation in response to physiological changes [38–40]. However, it remains uncertain whether GYS1 and NONO participate in transcriptional condensates through LLPS mechanisms to regulate MyoD-mediated myogenic programming and differentiation.

Herein, we report that nuclear GYS1 interacts with NONO and undergoes phase separation to form dynamic protein condensates that regulate enzyme export and glycogen synthesis. Furthermore, the nGYS1-NONO complex condenses with MyoD and the PIC to form transcriptional condensates, modulating C2C12 differentiation and cardiotoxin (CTX)-induced muscle regeneration in mice.

## Results

### GYS1 dynamically reassembles into subcellular protein bodies under conditions of glycogen depletion and transcriptional inhibition

GYS1 is a nucleocytoplasmic shuttling protein compartmentalized in the cytosol and nucleus of cells. At least three types of GYS1 protein bodies have been observed: (1) cytosolic GYS1 granules or glycosomes (of various sizes) under normal culture conditions; (2) nuclear GYS1 (nGYS1) puncta induced by the nuclear export inhibitor leptomycin B (LMB) or glucose deprivation; and (3) nGYS1 nucleolar caps induced by the transcription inhibitor actinomycin D (Act.D) (Fig. 1A, 1B; Fig. S1A, S1B; Table S1). nGYS1 puncta can be readily detected and were promoted by LMB in HeLa cells cultured in glucose-rich medium. In 293T cells, nGYS1 puncta were undetectable in glucose-rich culture but were significantly induced by glucose starvation, LMB, or Act.D treatment (Fig. 1B; Fig. S1B). Therefore, we used 293T cells to study the redistribution of GYS1 protein bodies under stress conditions. The transcription inhibitor Act.D, but not the translation inhibitor cycloheximide (CHX), induces the segregation of nGYS1 into the nucleolar cap (Fig. S1C), a typical structure of nucleoplasmic RBPs under transcriptional arrest [41]. The morphology of the GYS1 nucleolar cap was distinct from that of the cytosolic glycosomes or nGYS1 puncta induced by glucose starvation or LMB (Fig. 1B; Fig. S1B, S1C). Fractionation analysis revealed that the nuclear translocation of GYS1 was promoted by glucose starvation but not by Act.D (Fig. S1D). Meanwhile, Act.D did not affect glycogen levels or glycosome formation (Fig. 1B; Fig. S1E), suggesting that Act.D promotes the nucleolar concentration of nGYS1 rather than the nuclear translocation of cytosolic GYS1. The GYS1 nucleolar cap was specifically induced by the Pol II inhibitor DRB (5,6-dichlorobenzimidazole 1-β-D-ribofuranoside) but not by the Pol I inhibitor CX5461 or the Pol III inhibitor ML60218, although CX5461 appeared to enhance the effect of DRB (Fig. S1F, S1G).

**Fig. 1.**
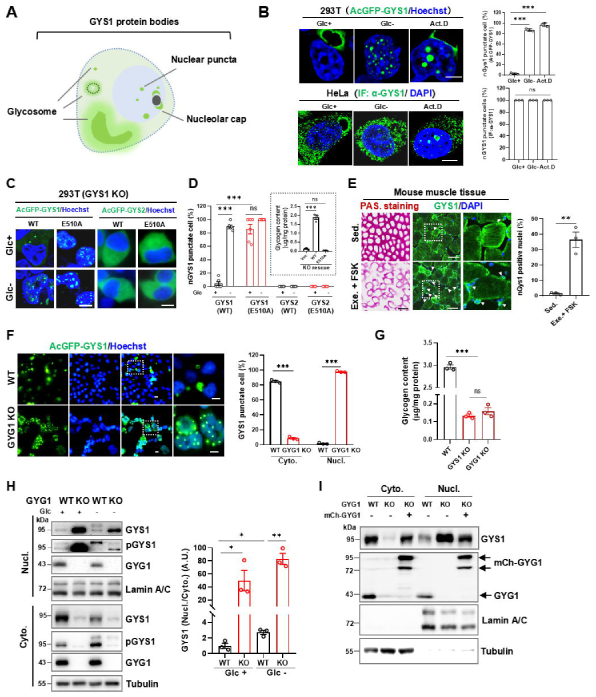
Reassembly of subcellular GYS1 protein bodies under energy or transcription stress conditions. **A** Diagram depicting three types of GYS1 protein bodies. **B** Subcellular distribution of GYS1 protein bodies. AcGFP-GYS1 in 293T cells or GYS1 IF staining in HeLa cells with the indicated treatment (left); nGYS1 punctate cells were quantified by counting the percentage of cells with more than five discernible fluorescent puncta in the nucleus (right). Cells were glucose-starved or treated with Act.D (1 μg/mL) for 12 h. **C** Imaging of AcGFP-GYS1/2 redistribution upon glucose starvation. GYS1-KO 293T cells were rescued with AcGFP-GYS1/2 WT or E510A mutants. **D** Quantification of nGYS1 punctate cells (>5 puncta per nucleus), and the inset shows the glycogen content of GYS1 WT- or E510A-rescued cells. **E** Nuclear accumulation of GYS1 in mouse myofibers. Muscle glycogen was depleted by intramuscular injection of 4 mg/kg FSK, followed by starvation for 12 h and then running to exhaustion; cross-sections of plantaris muscle were stained with PAS or IF. GYS1-positive nuclei were quantified in three sections for each group; scale bar, 100 μm. **F** Nuclear GYS1 puncta formation in GYG1-KO cells. AcGFP-GYS1 in normally cultured WT and *GYG1*-KO cells (left); GYS1 punctate cells in the cytosol and nucleus were quantified (right). **G** Glycogen contents in the WT, GYG1-, or GYS1-KO 293T cell lines. **H** IB analysis of cytosolic and nuclear GYS1 fractions in WT and GYG1-KO cells. nGYS1 was quantified by the signal ratio of cyto. and nucl. GYS1 (normalized to Lamin A/C and Tubulin, respectively), and then normalized to the first lane (=1.0); three experiments were performed. **I** IB analysis of subcellular GYS1 distribution in WT, GYG1-KO, and mCherry-GYG1-rescued cells. Error bars show the mean `±` SEM of 3 biological replicates; statistical analysis using two-tailed t-test; scale bars: 10 μm **(B, C, F)** and 100 μm **(E).**

To determine how the GYS1 protein body is spatially regulated from the cytosol to the nucleus, we reconstituted GYS1 mutants in GYS1 knockout (KO) 293T cells, excluding the oligomerization interference by endogenous GYS1 (Fig. S1A, S1H). Glycogen binding is essential for GYS1 compartmentalization in the glycosome, as the glycogen-binding-defect Y239A mutant [42] and the catalytically inactive E510A mutant [43] are no longer confined to the cytosol, even in the presence of glucose. Notably, the G6P-binding arginine-rich cluster mutant R1A/R2A, as well as the mutant with the simultaneous E510A mutation, was excluded from the nucleus, suggesting that the R1/R2 arginine cluster is critical for GYS1 nuclear import (Fig. S1H, S1I, and Table S2). Phosphorylations of GYS1 do not affect its translocation in response to glucose starvation, as previously reported [7, 8, 44]. Interestingly, GYS1`—`unlike its liver counterpart GYS2`—`forms nuclear puncta in response to glucose starvation. In contrast to GYS2 (E510A), the inactive GYS1 (E510A) forms nuclear puncta even in the presence of glucose (Fig. 1C, 1D). The nGYS1 protein bodies are unrelated to nuclear glycosomes, as no glycogen is synthesized in glucose-starved or E510A-rescued cells (Fig. 1D inset). Additionally, we observed an increase in nuclear translocation in mouse myofibers when glycogen was depleted by running exercise and treatment with the glycogenolysis agent forskolin (FSK) (Fig. 1E). To exclude the direct effect of glucose on the cytosolic retention of GYS1, we disrupted the glycogen initiator gene *GYG1* in 293T cells. Interestingly, most AcGFP-GYS1 in GYG1-KO cells accumulated as intranuclear bodies in glucose-rich medium, with no detectable glycosome GYS1 in the cytosol (Fig. 1F). The glycogen content of GYG1-KO cells decreased to a background level, similar to that in GYS1-KO 293T cells (Fig. 1G). Consistently, cell fractionation analysis revealed that GYS1 protein was mainly found in the nucleus of GYG1-KO cells, while rescue of GYG1 restored the cytosolic retention of GYS1 (Fig. 1H, 1I, Fig. S1J). These data suggest that glycogen, rather than glucose per se, is responsible for retaining GYS1 in the cytosolic glycosome.

## nGYS1 interacts and condenses with NONO in cells

To understand the regulation and function of nGYS1 protein bodies, we explored the nGYS1 interactome through tandem affinity purification (TAP) and live-cell TurboID proximity labeling [45]. In glucose-starved cells, the TurboID-GYS1 fusion enzyme efficiently biotinylated nuclear proteins, which were subsequently identified using mass spectrometry (MS) (Fig. S2A-S2E). We also purified nGYS1 using dithiobis (succinimidyl propionate) (DSP)-crosslinked TAP, followed by MS (TAP-MS) (Fig. S2D, S2F). Both methods revealed that nGYS1 is involved in RNA processing and transcriptional regulation (Fig. 2A, Fig. S2G-S2J). Interestingly, several paraspeckle proteins, including NONO, FUS, paraspeckle component 1 (PSPC1), splicing factor proline and glutamin rich (SFPQ), and RNA binding motif protein 14 (RBM14), were identified as nGYS1-interacting proteins (Fig. 2A), and these interactions were validated using TurboID-GYS1 labeling. In contrast, Lamin A/C and the nuclear speckle protein SC35 were not labeled and served as internal negative controls (Fig. S2K, S2L). Co-immunoprecipitation (Co-IP) analysis confirmed the interactions between endogenous NONO and GYS1 in 293T and HeLa cells under glucose starvation conditions (Fig. 2B, 2C; Fig. S3A). Additionally,

**Fig. 2.**
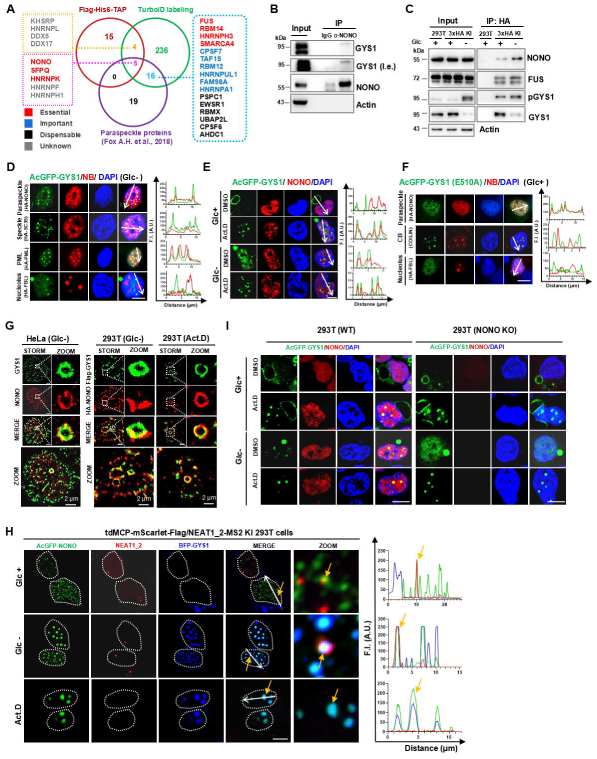
nGYS1 interacts with and condenses NONO in cells. **A** Venn diagram analysis of nGYS1-interacting RBPs identified by TurboID-MS, TAP-MS, and published paraspeckle proteins. **B** Coimmunoprecipitation (Co-IP) validation of the NONO-GYS1 interaction in glucose-starved 293T cells. **C** Co-IP analysis of the NONO-GYS1 interaction in GYS1^3xHA^ endogenously tagged 293T cells. **D** Colocalization of GYS1 and nuclear bodies. AcGFP-GYS1-expressing 293T cells were transfected with HA-tagged NB marker cDNA and IF-stained with HA antibody after glucose starvation. Colocalization was analyzed by the fluorescence intensity profile along a white arrow line crossing the nucleus. **E** Nuclear AcGFP-GYS1 and endogenous NONO colocalization in 293T cells with indicated treatments. **F** AcGFP-GYS1 (E510A) and NB colocalization in 293T cells cultured with glucose. **G** Dual-color STORM imaging of the nGYS1-NONO complex. Endogenous GYS1 and NONO in glucose-starved HeLa cells were costained with the indicated antibodies (left panel); Flag-GYS1 and HA-NONO co-expressing 293T cells were treated with glucose starvation or Act.D and costained with HA and Flag antibodies (right panel). **H** Colocalization of BFP-GYS1, AcGFP-NONO, and endogenous NEAT1_2 in mScarlet-labeled NEAT1_2 293T KI cells with the indicated treatments. Yellow arrows point to colocalized nGYS1/NONO (bottom), NONO/NEAT1_2 (top), or nGYS1/NONO/NEAT1_2 (middle). **I** Imaging of NONO or nGYS1 nucleolar caps induced by Act.D in WT and NONO-KO 293T cells. Scale bar: 10 μm.

NONO directly complexed with GYS1, independent of RNA bridging (Fig. S3B-S3F). Co-IP analysis mapped the GYS1 C-terminus (CT) as the direct binder of the RNA recognition motif (RRM) in NONO (Fig. S3C-S3F). Consistently, nGYS1 fully colocalized and formed nuclear body-like condensates with PSPC1, FUS, and SFPQ under glucose starvation conditions. It also partially colocalized with promyelocytic leukemia (PML) and overexpressed Cajal body protein COILIN. However, GYS1 did not colocalize with SC35, the nucleolar protein fibrillarin (FBL), or endogenous COILIN (Fig. 2D; Fig. S3G). Notably, when treated with Act.D, glucose starvation, or both, NONO consistently merged with nGYS1 as nuclear condensates (Fig. 2E; Fig. S3H). The GYS1 (E510A) mutant condensed with NONO under glucose-rich conditions but not with FBL or COILIN in the nucleus (Fig. 2F). Stochastic optical reconstruction microscopy (STORM) imaging showed that nGYS1 and NONO clustered together and merged into a ring structure (Fig. 2G). GYS1-CT mainly resides in the nucleus and forms granulated compartments with ectopic NONO independent of glucose; in contrast, GYS1 (`Δ`CT) is mostly excluded from the nucleus and no longer colocalizes with NONO (Fig. S3I). These data suggest that GYS1-CT, the NONO-binding domain, is crucial for nGYS1 condensation.

GYS1 was predicted to be an RBP with an interaction score of 0.65 (Fig. S3J). The paraspeckle lncRNA NEAT1_2 was identified as the top candidate binding to GYS1 using catRAPID online tools [46], which was validated by RNA immunoprecipitation-qPCR (Fig. S3K, S3L). Consistently, nGYS1 and NONO colocalized with paraspeckles in mScarlet-labeled Neat1_2 knock-in cells [47] (Fig. 2H). The nGYS1-NONO colocalization was maintained even when NEAT1_2 was abolished by Act.D treatment (Fig. 2H, 2I; Fig. S1G), suggesting that NEAT1_2 or paraspeckles are not essential for GYS1-NONO complex formation, which is consistent with the biochemical data (Fig. S3B-S3F).

## NONO retains GYS1 in the nucleus to restrain glycogenesis and support strenuous exercise in mice

We found that NONO is essential for the effective formation of nGYS1 protein bodies induced by glucose starvation. In NONO-KO cells, the percentage of nGYS1 punctate cells was significantly reduced; however, this reduction can be restored by reintroducing NONO (Fig. 3A, 3B). Notably, NONO is not required for the formation of nGYS1 nucleolar caps induced by Act. D (Fig. 2I). The glycogen content in NONO-KO cells increased approximately threefold compared to wild-type (WT) cells but decreased after NONO rescue (Fig. 3C, 3D). No change in GYS1 phosphorylation was detected in NONO-KO cells (Fig. S4A); in contrast, nGYS1 protein levels were higher in WT and NONO-rescued cells than in KO cells under both normal culture and glucose restimulation conditions (Fig. S4B), suggesting that NONO restricts nGYS1 export. Indeed, cytosolic glycosomes accumulated more rapidly upon glucose restimulation in NONO-KO cells than in NONO-rescued cells (Fig. S4C). The data presented above (Fig. 1-Fig. 3) support a model in which GYS1 nucleocytoplasmic shuttling is balanced by cytosolic glycosome binding and nuclear condensate retention. Glycogen depletion or loss of NONO disrupts subpool homeostasis, inducing GYS1 translocation and subcellular protein body reorganization (Fig. 3E).

**Fig. 3.**
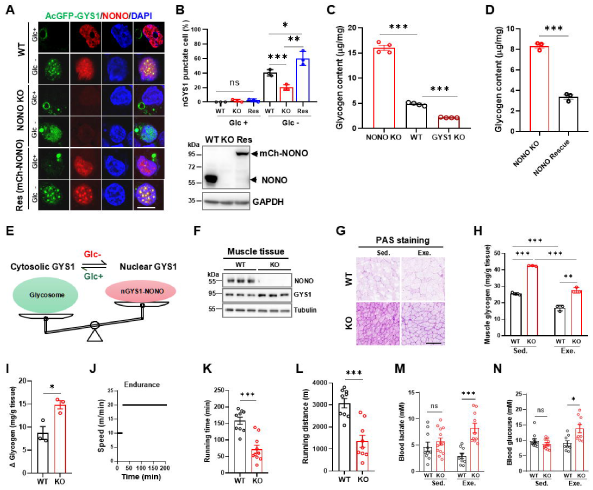
NONO condenses GYS1 in the nucleus to inhibit glycogenesis and support strenuous exercise in mice. **A** Nuclear AcGFP-GYS1 puncta formation in WT, NONO-KO, and mCherry-NONO-rescued 293T cells under the indicated conditions. **B** IB analysis of NONO expression (top) and quantification of nGYS1 punctate cells in WT, NONO-KO, and mCherry-NONO-rescued cells. **C** Glycogen content in WT and KO cells, with two clonal cell lines shown for each genotype. **D** Glycogen content in NONO-KO and mCherry-NONO-rescued 293T cells. **E** A graphic model illustrating that the cytosolic and nuclear GYS1 subpools are balanced by cytosolic glycosome binding and nuclear condensates retention. **F-I** Glycogen contents in sedentary (Sed.) or 70-min exercised (Exe.) muscle tissues of WT and Nono-KO mice. IB analysis of Nono and Gys1 in WT and KO mice **(F);** Glycogen PAS staining in muscle **(G)**; glycogen content of skeletal muscle tissues **(H)**; and consumption of muscle glycogen during exercise **(I)**; n = 3 mice per group. **J-L** Exercise endurance test in WT and Nono-KO mice. A diagram depicting increments of speed over time **(J)**, running time **(K)**, and running distance **(L)** for the motorized treadmill test; n = 9-10 mice per group. **M, N** Blood lactate **(M)** and blood glucose levels **(N)** in sedentary or 70-min exercised mice; n = 8-13 mice per group. Error bars show the mean `±` SEM of 3 biological replicates for each group; statistical analysis using two-tailed t-test; scale bars: 10 μm **(A)** and 100 μm **(F)**.

Muscle glycogen levels are associated with strenuous exercise performance [48]. We detected higher glycogen levels in the muscles of Nono-KO mice compared to their WT littermates under both sedentary and exercised conditions (Fig. 3F-3H). However, glycogen consumption in the muscles of Nono-KO mice was approximately 50% higher than in WT control mice (Fig. 3I). An assessment of acute running endurance performance on a motorized treadmill showed that Nono-KO mice ran for shorter times and distances (about 30% less) than their WT controls (Fig. 3J-3L). Post-exercise blood glucose and lactate levels were elevated (Fig. 3M, 3N), while blood triglyceride and nonesterified fatty acid levels remained normal in Nono-KO mice (Fig. S4D, S4E). These data suggest that Nono deficiency promotes glycolytic metabolism due to excessive glycogen deposition in muscle, a phenotype similar to Gyg-KO mice [49].

### GYS1 undergoes protein phase separation and condenses with NONO or FUS *in vitro*

Protein phase separation (PPS) has been proposed as a spatiotemporal organizing principle for metabolic enzymes that actively regulate enzyme functions, including biochemical reaction integration and gene expression. NONO and FUS, both of which interact and condense with nGYS1 (Fig. 2A, 2C; Fig. S2K, S2L, S3G), are known as TFs and RBPs with LLPS properties [26–28]. The termini of both GYS1 and NONO, as predicted by Fuzdrop [50], contain droplet-promoting regions (DPRs) (Fig. 4A). To test whether GYS1 undergoes PPS *in vitro*, we purified flag-tagged AcGFP-GYS1, mCherry-NONO, and mCherry-FUS proteins from 293T cells (Fig. S5A). AcGFP-GYS1 was co-purified with ectopic GYG1 (Y195F), which increased the solubility of GYS1 [51, 52]. Purified GYS1 formed micrometer-sized spheres *in vitro* and coalesced into larger droplets over time (Fig. 4B). However, the fluorescence recovery after photobleaching (FRAP) assay revealed that the GYS1 spheres were not dynamic (Fig. 4C). In contrast, mCherry-NONO formed highly mobile, liquid-like condensates (Fig. 4D, 4E). When GYS1 and NONO were mixed, they co-condensed as liquid coacervates, with both proteins exhibiting high mobility (Fig. 4F, 4G). Similar results were obtained for GYS1 and FUS (Fig. S5B, S5C). Moreover, GYS1 formed larger expanded droplets when mixed with NONO or FUS (Fig. S5D). AcGFP-GYG1(Y195F) alone did not form droplets, and AcGFP-GYS1 purified from GYG1-KO cells exhibited the same ability to form spheres, albeit with a low purification yield (Fig. S5E), indicating that GYS1 forms droplets independently of GYG1. The formation of GYS1 spheres was inhibited when mixed with the LLPS inhibitor 1,6-hexanediol (HEX), which disrupts weak hydrophobic interactions. However, it was not affected by RNase A or amylase treatment (Fig. S5F), suggesting that RNA or glycogen is unnecessary for GYS1 PPS *in vitro*. GYS1 forms droplets across a wide range of salt concentrations. Notably, the addition of the crowding agent PEG8000, bulk RNA, or genomic DNA strongly promoted GYS1 droplet formation (Fig. 4H, Fig. S5G-S5I), and the GYS1-RNA droplets were also immobile gel-like structures (Fig. 4I). Purified liver glycogen undergoes phase separation without the involvement of proteins [23]. Indeed, both glycogen and GYS1 undergo phase separation *in vitro*; GYS1 further merged into glycogen droplets after mixing the two phases, and the GYS1-glycogen droplets were not dynamic (Fig. S5J-S5L). These data suggest that GYS1-RNA and GYS1-glycogen droplets are biophysically distinct from GYS1-NONO and GYS1-FUS condensates.

**Fig. 4.**
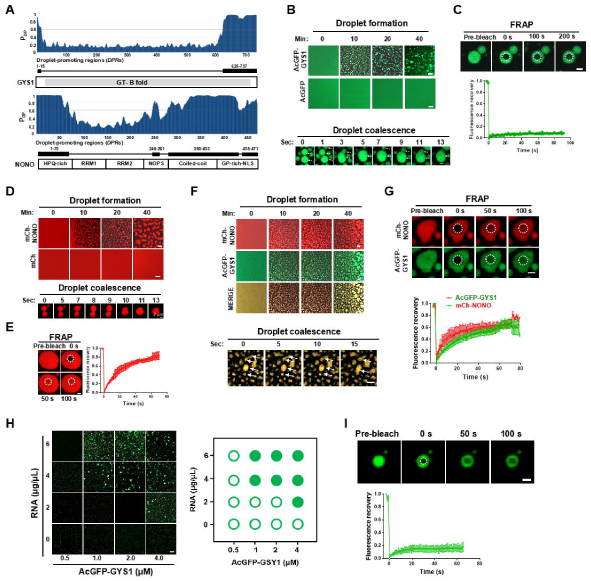
GYS1 protein phase separates and co-condensates with NONO *in vitro.* **A** Residue-based droplet-promoting probability (P_DP_) analysis of structured GYS1 and NONO using the Fuzdrop algorithm. **B** Droplet formation for 4 μM purified AcGFP or AcGFP-GYS1 in buffer containing 50 mM Tris-HCl (pH 7.5), 150 mM NaCl at 25 °C with the indicated incubation time (top), and time-lapse images of droplet coalescence (bottom); scale bars, 2 µm. **C** FRAP assay of AcGFP-GYS1 droplets. Representative time-series images show the fluorescence recovery of the AcGFP-GYS1 droplet after photobleaching (top). Quantified FRAP signals from n = 7 droplets (bottom); scale bars, 2 µm. **D** Protein phase separation of 3 μM purified mCherry-NONO with the indicated incubation time (top) and time-lapse images of droplet coalescence (bottom); scale bars, 2 µm. **E** FRAP assay of mCherry-NONO droplets (left) and quantified FRAP signals from n = 3 droplets (right); scale bars, 2 µm. **F** Co-condensation of mCherry-NONO (3 μM) and AcGFP-GYS1 (3 μM) mixtures at the indicated times (top) and time-lapse images of droplet coalescence (bottom); scale bars, 2 µm. **G** FRAP assay on droplets of the mCherry-NONO and AcGFP-GYS1 mixture (top) and quantified FRAP signals from n = 8 droplets (bottom) in the green and red channels; scale bars, 2 µm. **H** Representative images of GYS1-RNA droplets 5 min after mixing varying concentrations of protein and RNA (left) and a corresponding phase diagram (right). Filled circle, droplet formation; empty circle, no droplet formation. **I** FRAP assay on droplets of RNA (600 ng/μL) and AcGFP-GYS1 (4 μM) mixture (top) and quantification of the FRAP assay for n = 6 droplets (bottom); scale bars, 2 µm. Error bars represent the mean `±` SEM; scale bar, 10 μm unless otherwise specified.

## NONO spatiotemporally regulates nGYS1 protein body dynamics

To further characterize the mobility of GYS1 protein bodies at distinct subcellular locations, we compared the dynamics of glycosome GYS1 under normal culture conditions with the nGYS1-NONO condensates induced by glucose starvation. The nGYS1 is highly dynamic, similar to the colocalizing mCh-NONO, whereas the glycosome GYS1, spatially separated from nuclear NONO, is an immobile structure (Fig. S6A). This finding is consistent with *in vitro* phase separation data. Differences in the mobility of distinct subcellular GYS1 protein bodies were also observed in the same cell after short-term glucose restimulation, which led to the coexistence of nuclear and cytosolic AcGFP-GYS1 protein bodies (Fig. 5A). Notably, the nucleus-localized GYS1 (E510A) protein bodies were highly mobile, independent of glucose (Fig. 5B). However, both the phosphomimetic mutant 9SD and the nonphosphorylatable mutant 9SA protein bodies were immobile in the cytosol and nucleus (Fig. S6B). In GYS1-KO HeLa cells expressing AcGFP-GYS1 at the endogenous level (AcGFP-GYS1^endol^), nuclear puncta were induced by various treatments (Fig. S6C, S6D). We compared the mobility of cytosolic GYS1 and nuclear GYS1 puncta, which coexist in FSK-treated AcGFP-GYS1^endol^ HeLa cells. Consistently, the nGYS1 puncta were more dynamic than the cytosolic puncta (Fig. 5C). Additionally, subcellularly distinct GYS1 protein bodies showed different sensitivities to HEX. Most nGYS1 protein bodies were dissolved by 5% HEX treatment at 300 seconds (s), while more than 60% of the cytosolic GYS1 microbodies remained (Fig. 5D, 5E). NONO overexpression promoted HEX-induced nGYS1 microbody disassembly within 120 s (Fig. S6E). Notably, the GYS1 (E510A) condensates in the nucleus were mostly dissolved within 30 s (Fig. 5D, 5F); in contrast, glycosome GYS1 could only be dissolved by higher concentrations (10% or 15%) of HEX (Fig. S6F). These data suggest that the dynamics of GYS1 protein bodies are spatiotemporally regulated by distinct subcellular partners, specifically glycogen and NONO.

**Fig. 5.**
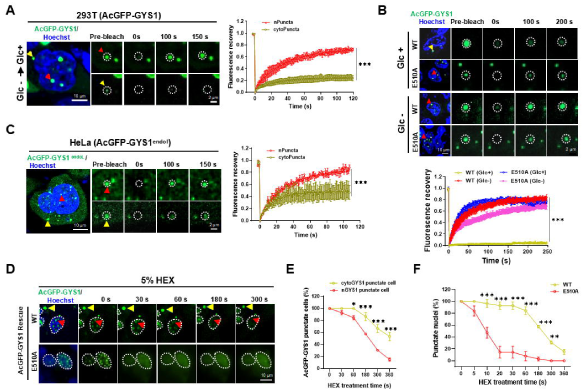
The nGYS1 condensates are more dynamic than the glycosome GYS1. **A** Representative images and FRAP assay for ectopic AcGFP-GYS1 nuclear and cytosolic puncta in 293T cells restimulated with 25 mM glucose for 5 h after 12 h of glucose deprivation (left) and quantified FRAP signals for each type of puncta (n = 5) (right). **B** Representative images and FRAP assay for ectopic AcGFP-GYS1 WT or E510A nuclear and cytosolic puncta in 293T GYS1-KO cells treated with or without glucose starvation (top) and quantification of the FRAP signals for each type of puncta (n =5) (bottom). **C** Representative images and FRAP assay for endogenous-level AcGFP-GYS1 nuclear puncta (red arrowhead) and cytosolic puncta (yellow arrowhead) in HeLa cells treated with FSK (100 μM, 6 h) (left) and quantification of the FRAP signals for each type of puncta (n = 3) (right). **D-F** Diffusion of AcGFP-GYS1 WT or E510A puncta in 293T GYS1-KO cells. Representative images of AcGFP-GYS1 WT or E510A puncta after 5% HEX treatment for the indicated times **(D. T**he nuclear and cytosolic punctate cells in AcGFP-GYS1-transfected cells were counted in three fields of view (FOVs) and normalized to zero time point (=100%) **(E)**. The punctate nuclei of AcGFP-GYS1 (WT) and (E510A) were plotted by counting the punctate nuclei in three FOVs for each HEX treatment **(F)**; a punctate cell was defined as more than 5 puncta in the nucleus or cytosol. Error bars show the mean `±` SEM. **A, C** were statistically analyzed by unpaired two-tailed t-test; **B** by one-way ANOVA; **E, F** by multiple t-test. Scale bars are labeled in each image.

### The nGys1-Nono complex regulates C2C12 differentiation and myofiber development

Dysregulation of GYS1 and glycogen metabolism leads to cardiomyopathy and exercise intolerance in affected GSD patients [53]. 80% of male Nono-KO and 90% of Gys1-KO mice die from heart development defects before birth [54, 55]. Compared to their WT littermates, the surviving Nono-KO mice exhibited smaller body size and decreased body weight (Fig. 6A, 6B). The gastrocnemius (GC) and tibialis anterior (TA) muscles of the Nono-KO mice exhibited a decrease in size, weight, and muscle/body weight ratio compared to the WT controls (Fig. 6C-6F), indicating defects in myofiber development. Indeed, the average cross-sectional area (CSA) of the myofibers in the KO mice decreased, as determined by wheat germ agglutinin (WGA) staining (Fig. 6G). Consequently, more small fibers and fewer large fibers were observed in the KO mice than in the WT controls (Fig. 6H). No change in the proportion of type I or type II fibers in GC muscle was detected in KO mice, as indicated by immunostaining for myosin heavy chain 1 (MHC1) and MHC2b (Fig. S7A). These data suggest that Nono is important for myofiber development in mice. Using the C2C12 cell model of myogenesis, we found that both Gys1 and Nono proteins increased during C2C12 differentiation, coinciding with the expression of the differentiation markers MyHC and MyoD (Fig. 6I). The level of paraspeckle Neat1_2 RNA also increased alongside MyHC and Myogenin, as previously reported [56] (Fig. S7B). Gys1 prominently colocalized with Nono in glucose-containing differentiation medium, and this colocalization was further enhanced by FSK treatment (Fig. S7C). Pretreatment with glucose deprivation or FSK, both of which induce nGYS1 puncta, promoted C2C12 differentiation, while Act.D or HEX completely prevented it (Fig. S7D). These data indicate that nGYS1 puncta promotes C2C12 differentiation. Indeed, Gys1- or Nono-KO C2C12 cells failed to differentiate (Fig. 6J-6M). In contrast, C2C12 cells deficient in the lncRNA Neat1-2, the primary scaffold for paraspeckles, differentiated more efficiently than WT cells (Fig. S7E-S7I). However, C2C12 cells with Nono and Neat1-2 double KO (DKO) or Gys1 and Neat1-2 DKO failed to differentiate (Fig. S7F-S7I). These data suggest that Gys1 and Nono, but not paraspeckles, are essential for C2C12 differentiation. GYS1 rescue experiments showed that the R1A/R2A mutant, which mainly resides in the cytosol, was ineffective at rescuing Gys1-KO cell differentiation. In contrast, punctate GYS1 (E510A) and GYS1-CT partially rescued differentiation (Fig. 6N, 6O). Additionally, the RRM domain of NONO, which is the GYS1 binding region, partially rescued Nono-KO C2C12 cell differentiation, and deletion of the RRM largely compromised the rescue effect (Fig. 6P). These data suggest that the nGys1-Nono condensates are crucial for myogenic differentiation.

**Fig. 6.**
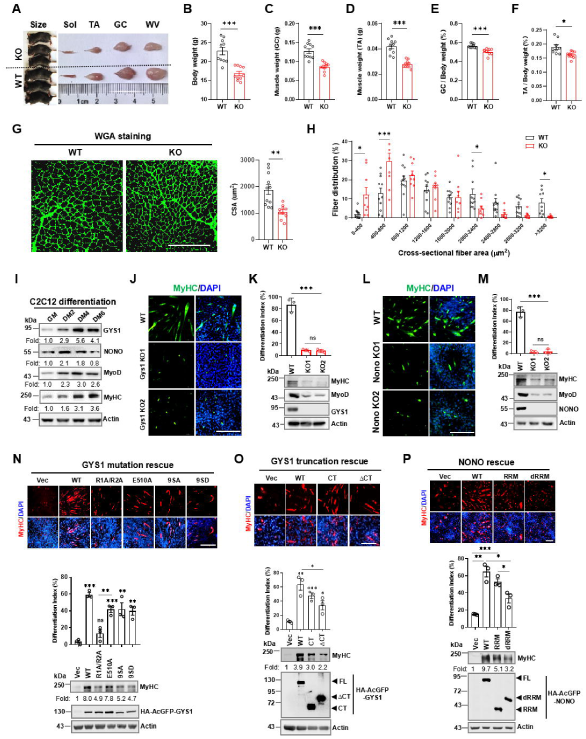
Nono deficiency leads to myogenic defects in mouse skeletal muscle. **A** Body and muscle size of WT and Nono-KO mice**. B-F** Body and muscle weights of WT and Nono-KO mice. Body weights of WT and Nono-KO mice **(B)**, n = 10 mice per group. Total GC **(C)** or TA **(D)** muscle weight and the GC muscle/body weight ratio **(E)** or TA muscle/body weight ratio **(F)** for WT and Nono-KO mice; n = 9-10 mice per group. **G, H** WGA staining (left) and average cross-sectional area (CSA) (right) of the GC myofibers **(G)** and the frequency distribution of myofiber CSA in WT and Nono-KO mice **(H)**; statistical analysis using unpaired t-test; n = 9-11 mice per group. **I** IB analysis of protein levels by indicated antibodies during C2C12 differentiation. **J-M** Differentiation of Gys1- or Nono-KO C2C12 myoblasts. Representative images of MyHC staining in WT, GYS1-KO **(J)**, or Nono-KO **(L)** C2C12 cells after 4 days of differentiation. The differentiation index was calculated by the ratio of the number of nuclei in the myotubes to the total number of nuclei in one FOV **(K, M top)**, and MyHC protein levels in C2C12 myoblasts with the indicated genotypes **(K, M, bottom)** were analyzed. **N-P** Differentiation of GYS1-KO C2C12 myoblasts rescued by GYS1 point mutants **(N)** or truncations **(O)**; MyHC IF staining (top) and differentiation index (middle) are shown with the expression of the rescued GYS1 mutants (bottom) after 4 days of differentiation. Nono-KO C2C12 cells were rescued by NONO RRM truncation or RRM deletion **(P)**. MyHC staining (top) and quantified differentiation index (middle) were shown with the expression of the rescued NONO mutants (bottom) after 4-day differentiation. Error bars show the mean `±` SEM; statistical analysis using unpaired t-test; scale bars: 100 μm **(G)** and 50 μm **(J, L, N, O, P)**

### nGYS1-NONO co-condenses with MyoD and the pre-initiation complex (PIC) to regulate myogenic gene expression and cardiotoxin (CTX)-induced muscle regeneration in mice

To understand how nGYS1-NONO puncta regulate myogenesis, we tested whether GYS1-NONO forms complexes with MyoD to modulate myogenic reprogramming. Indeed, GYS1, NONO, and MyoD formed complexes as nuclear condensates in C2C12 myoblasts cultured in differentiation medium (DM) or treated with FSK (Fig. 7A, 7B; Fig. S8A). nGYS1 did not colocalize with SC35 or MyHC (Fig. S8B, S8C). NONO and MyoD co-condensed with nGYS1 without significant differences among the WT, 9SA, and 9SD mutants (Fig. S8D). Furthermore, purified BFP-MyoD formed droplets with GYS1, NONO, or both, while MyoD alone tended to aggregate *in vitro* (Fig. 7C). Next, we investigated whether the GYS1-NONO-MyoD complex is part of the transcriptional condensates that regulate transcription initiation, as proposed in a recent model of gene expression regulation [57, 58]. Indeed, NONO, GYS1, and active Pol II, indicated by C-terminal Ser-5 phosphorylation (pS-5), were detected in the MyoD Co-IPs. Multiple-component interactions and colocalizations were enhanced when the culture was switched from growth medium (GM) to DM (Fig. 7D, Fig. S8E). Gys1 or Nono deficiency reduced the MyoD-Pol II (pS-5) interaction (Fig. 7E). The rescue of the RRM-deleted mutant of NONO largely diminished the co-condensation of nGYS1-NONO-MyoD, as well as the nGYS1 condensates in Nono-KO C2C12 cells (Fig. S8F). Furthermore, nGYS1 was found to interact and co-condense with major PIC components, including BRD4, TBP, GTF2H5, and MED28 (Fig. S8G-S8M), through Co-IP, colocalization, and TurboID-GYS1 proximity labeling. The condensation of the GYS1-NONO complex and Pol II (pS-5) appears to depend on RNA transcription, as their colocalization was weakened by Act.D treatment (Fig. S8N). Consistently, the mRNA levels of MyoD target genes, including *Myogenin*, *Tnni2*, and *Myomixer,* were significantly down-regulated in Gys1 and Nono-KO C2C12 cells during differentiation (Fig. 7F). MyoD is a basic helix-loop-helix (bHLH) TF that binds to E-box elements of myogenic gene promoters to mediate transcription. The absence of either Gys1 or Nono leads to a decrease in MyoD occupancy on the E-box-containing promoter regions of the *Myogenin*, *Tnni2*, and *Myomixer* genes (Fig. 7G). Therefore, multivalent interactions among GYS1, NONO, MyoD, PIC components, and nascent RNA are involved in the assembly of transcriptional condensates that drive the myogenic gene transcription.

**Fig. 7.**
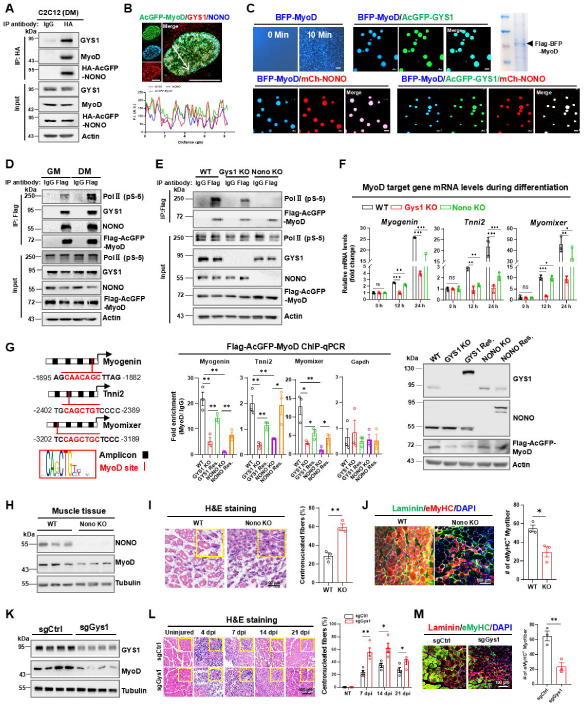
GYS1-NONO co-condenses with MyoD and Pol II to regulate myogenic gene transcription. **A** Co-IP analysis of NONO, GYS1, and MyoD interactions in C2C12 myoblasts cultured in glucose-free DM for 24 h. **B** Colocalization of AcGFP-MyoD, Nono, and Gys1 proteins in C2C12 cells cultured in DM for 24 h. **C** *In vitro* droplet formation of BFP-MyoD (1 μM), AcGFP-GYS1 (4 μM), mCherry-NONO (2 μM), or all three mixed. **D** Co-IP analysis of the interactions between MyoD and Nono, Gys1, or Pol II (pS-5) in C2C12 myoblasts cultured in GM or DM for 24 h. **E** MyoD-Pol II (pS-5) interaction in C2C12 WT and the indicated KO cells cultured in DM. **F** MyoD targets mRNA levels in the indicated C2C12 myoblasts cultured in DM. **G** A schematic depicting the E-box regions of target gene promoters (left), and ChIP-qPCR analysis of MyoD-occupied amplicon of target genes in various C2C12 myoblast cell lines cultured in DM (right). **H** IB analysis of muscle tissues from WT and Nono-KO mice. **I** H&E staining of TA muscle cross-sections (left) and quantification of centronucleated myofibers 7dpi of CTX (right); 200 myofibers/mouse. **J** Laminin and eMyHC immunostaining of TA muscles at 7 dpi (left), and the percentage of eMyHC**^+^** myofibers was quantified for Laminin and eMyHC-stained myofibers in three FOVs/section (right); n = 3-4 mice per group. **K** IB analysis of Gys1 in muscle tissues 28 days after AAV9 injection. **L** H&E staining of TA muscle cross-sections from AAV9-infected muscles, and the centronucleated myofibers were quantified (right). **M** Laminin and eMyHC immunostaining of TA muscles at 7 dpi (left), and the eMyHC**^+^** myofibers were quantified in three FOVs/section (right); n = 5-6 mice per group. Error bars show the mean ± SEM; statistical analysis using unpaired t-test; scale bars: 10 μm **(B)** and 100 μm **( I, J, L, M).**

To investigate the physiological roles of Gys1 and Nono in muscle regeneration, we first generated an acute skeletal muscle regeneration model using a single intramuscular injection of CTX in WT and Nono-KO mice (Fig. S9A). Compared to WT control muscle, Nono deficiency resulted in impaired muscle regeneration induced by CTX, as indicated by lower MyoD protein levels and a higher percentage of centronucleated myofibers (Fig. 7H, 7I). Additionally, embryonic myosin heavy chain (eMyHC) staining, an indicator of regenerating myofibers, also decreased in Nono-KO muscles (Fig. 7J). We generated Gys1- or Nono-deficient muscle by delivering sgRNA via recombinant adeno-associated virus 9 (AAV9) into the TA muscles. Cre expression, driven by the muscle hybrid (MH) promoter [59], induced muscle-specific expression of Cas9 and gene knockout in Rosa26-CAG-LSL-Cas9 knockin mice. The coexpression of NLuc with Cre marked the AAV9-infected area in live mice (Fig. S9A, S9B). Loss of Gys1 in muscle tissue led to impaired regenerative capability, as indicated by decreased MyoD levels, a higher percentage of centronucleated myofibers, and decreased eMyHC staining in Gys1-deficient muscle (Fig. 7K-7M). Similar regenerative defects were also observed in Nono-deficient muscle (Fig. S9C-S9E). Collectively, these results demonstrate that the Gys1-Nono complex is an important regulator of muscle regeneration in CTX-induced muscle regeneration in mice.

## Discussion

Eukaryotic metabolism is a highly orchestrated process involving various dynamic enzyme compartments that fluctuate with physiological rhythms. Stress-induced subcellular compartmentalization of metabolic enzymes has been frequently observed, representing a general model of regulation. GYS1 diffuses and translocates into the nucleus to form punctate protein bodies when glycogen is depleted, as demonstrated in glucose-starved and GYG1-KO cells (Fig. 1B, 1F). Thus, GYS1 senses cellular glycogen levels or energy status via glycosome formation and translocation. To better understand the regulation and transition of protein bodies in response to metabolic changes, we identified several TFs and RBPs that interact with GYS1 and compartmentalize together as nuclear condensates (Fig. 2A, 2D, and Fig. S2H). Among these, NONO condenses with nGYS1 to form puncta or nucleolar caps under conditions of glycogen depletion or transcription arrest, respectively (Fig. 2D, 2E). These condensates maintain the GYS1 nuclear subpool and restrict its export during glucose-stimulated glycogenesis. NONO deficiency reduces nGYS1 condensation and induces glycosome formation by shifting the GYS1 balance toward the cytosolic subpool (Fig. 3A-3E, Fig. S4B, S4C). Notably, both phosphomimetic and nonphosphorylatable GYS1 shuttled normally in response to glucose (Fig. S1H, S1I) and condensed with NONO and MyoD in C2C12 cells (Fig. S8D). Therefore, nGYS1 accumulation is not directly regulated by GYS1 phosphorylation but is more likely a secondary effect of GYS1 phospho-inactivation, which induces glycogen depletion and GYS1 nuclear condensation. Collectively, these findings support the model that GYS1 nucleocytoplasmic shuttling is balanced by cytosolic glycosome binding and the retention of its nuclear condensates.

The Rf motif of metabolic enzymes has been proposed to function as an RNA interaction domain [12, 60–64]. GYS1 contains two Rf domains and has been reported to bind various RNA species [13, 14]. In this study, GYS1 was found to colocalize and interact with lncRNA NEAT1_2 (Fig. 2H and Fig. S3J-S3L), suggesting that nGYS1 partially exists as a ribonucleoprotein. Paraspeckles are RBP-centered nuclear bodies scaffolded by lncRNA NEAT1_2, with their number increasing during C2C12 differentiation [56]. However, NEAT1_2_is not required for the GYS1-NONO interaction. Both NONO and GYS1 promote C2C12 differentiation, while Neat1_2 inhibits it (Fig. 2H, 2I, Fig. S3B-S3F, and Fig. S7E-S7I). Therefore, the Gys1-Nono complex acts independently of Neat1_2 or paraspeckles to modulate myogenesis. Nonetheless, the physiological role of paraspeckle-localized GYS1 requires further investigation. nGYS1 also condenses with other RBPs, such as FUS, which may link GYS1 to neurodegenerative diseases [65, 66]. Precise studies dissecting the nGYS1 interactome, or the binding RNA in nuclear bodies, would be helpful for fully defining the composition and function of various nGYS1 compartments.

Interestingly, purified GYS1 forms gel-like spheres or droplets *in vitro* and less mobile cytosolic glycosomes in cells, as evidenced by FRAP experiments (Fig. 4B, 4C, 5A, 5C). However, the GYS1 protein bodies undergo dynamic condensation when complexed with NONO or FUS, but not with glycogen or RNA *in vitro* (Fig. 4F-4I, Fig. S5C, S5L), highlighting the differential regulation of GYS1’s biophysical properties by distinct binding partners. *In vitro* droplet assays showed that GYS1 and NONO form droplets individually. MyoD alone tended to aggregate and form droplets only with GYS1 or NONO (Fig. 4B, 4D, Fig. 7C). Nuclear body reorganization has been reported to regulate myogenic gene transcription [67]; indeed, the deficiency of either Gys1 or Nono in C2C12 myoblasts blocks myogenic differentiation (Fig. 6J-6M). Consistently, mice deficient in Gys1 or Nono exhibited compromised muscle regeneration after CTX-induced acute injury (Fig. 7H-7M, Fig. S9C-S9E). Our mechanistic studies revealed that nGYS1-NONO interacts and co-condenses with MyoD and major PIC components, which includes Pol II (p-S5), BRD4, MED28, GTF2H5, and TBP, forming transcriptional condensates that drive myogenic gene expression (Fig. 7A-7E, Fig. S8A-S8M). A deficiency in either Gys1 or Nono compromised the interaction between MyoD and active Pol II, inhibiting myogenic gene expression (Fig. 7E-7G). Depletion of glycogen through pretreatment with glucose starvation or FSK induces nGYS1 condensates and promotes C2C12 differentiation (Fig. S7D). The rescue of nuclear punctate GYS1-CT and the E510A mutant in GYS1-KO cells, as well as NONO-RRM rescue in Nono-KO cells, partially restored C2C12 differentiation (Fig. 6N-6P). Collectively, these data suggest that nGYS1-NONO puncta are part of transcriptional condensates and are essential for MyoD-mediated myoblast differentiation. Nono or Gys1 deficiency is partially lethal due to heart development defects in mice. We found that the surviving Nono-KO mice exhibited an exercise intolerance phenotype similar to that of Gyg-KO mice, characterized by elevated glycogen accumulation in muscle, decreased running capability, and a phenotype of glycolytic energy expenditure [49] (Fig. 3F-3N). These data suggest that Nono and Gys1 play roles in both metabolic regulation and myogenic differentiation in muscle physiology, potentially linking exercise-induced glycogen depletion to muscle growth.

As summarized in Fig. 8, this study demonstrated that the condensation of nGYS1 and NONO through protein phase separation spatiotemporally regulates glycogenesis and MyoD-mediated myogenic transcription, providing mechanistic insights into the role of nuclear GYS1 in metabolic disorders and muscular dystrophy.

**Fig. 8.**
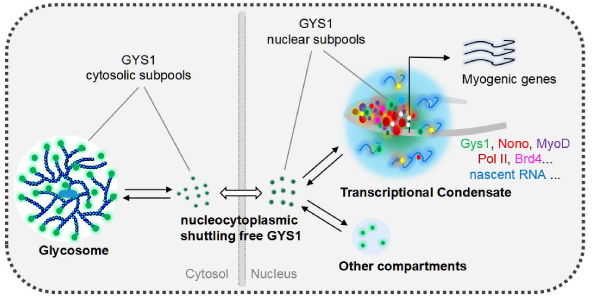
A diagram depicting the spatiotemporal regulation and subcellular function of GYS1. Two subpools of GYS1 are delicately equilibrated between the cytosolic glycosomes and nGYS1 condensates, depending on the cellular energy or physiological state. Among the various nGYS1 compartments, nGYS1-NONO condenses with MyoD to form transcriptional condensates centered on the PIC, regulating myogenic gene transcription and subsequent myoblast differentiation or muscle regeneration.

## List of Abbreviations

GSDs: glycogen storage diseases
Rf: rossmann-fold
G6P: glucose-6-phosphate
RBP: RNA-binding protein
NEAT1_2: nuclear paraspeckle assembly transcript 1 (NEAT1) isoform 2
DBHS: Drosophila behavior and human splicing
FUS: fused in sarcoma
RBM14: RNA binding motif protein 14
PSPC1: paraspeckle component 11
PML: promyelocytic leukemia
SC35: splicing component 35K
FBL: fibrillarin
SFPQ: splicing factor proline and glutamine rich
Act.D: actinomycin D
LMB: leptomycin B
CHX: cycloheximide
FSK: forskolin
HEX: 1,6-hexanediol
Myod1 or MyoD: myogenic differentiation 1
Myog: myogenin
MyHC: myosin heavy chain
Pol II: RNA polymerase II
DRB: 5,6-dichlorobenzimidazole 1-β-D-ribofuranoside
GC: gastrocnemius
TA: tibialis anterior
LLPS: liquid_Jliquid phase separation
PPS: protein-phase separation
MLOs: membrane-less organelles
IDRs: intrinsically disordered regions
NB: nuclear body
RRM: RNA recognition motif
Co-IP: co-immunoprecipitation
TAP: tandem affinity purification
STORM: stochastic optical reconstruction microscopy
FRAP: fluorescence recovery after photobleaching.

## Acknowledgments

We thank the Core Facilities at the State Key Laboratory of Oncology in South China and the Medical Science Public Platform of Shenzhen Campus, Sun Yat-sen University, for their assistance with imaging experiments. The mScarlet-labeled MS2-tagged Neat1_2 knock-in cell line was provided by Professor Zhou Songyang from Sun Yat-sen University. The GYS2 and SC35 cDNAs were provided by Professor Ronggui Hu at the Shanghai Institute of Biochemistry and Cell Biology, while the AAV vectors were provided by Professor Qiurong Ding from the Shanghai Institute of Nutrition and Health.

## Author contributions

X. Xie conceived the idea; S. Peng, C. Li, Y. Wang, Y. Yi, X. Chen, Y. Yin, Z. Gan, and X. Xie designed and performed the experiments, with contributions from F. Yang, F. Chen, Y. Ouyang, H. Xu, B. Chen, H. Shi, Y. Zhao, and L. Feng; Q. Li performed TurboID MS analysis. S. Peng, C. Li, Y. Wang, Y. Yi, X. Chen, L. Feng, Z. Gan, and X. Xie analyzed the data and wrote the manuscript with input from all other authors. L. Feng, Z. Gan, and X. Xie supervised the project.

## Data availability

All data necessary to evaluate the conclusions of this study are presented in the paper and/or the supplementary information. The mass spectrometry proteomics data have been deposited in the ProteomeXchange Consortium via the PRIDE partner repository, with dataset identifiers PXD051836 and PXD058561.

## Materials and methods

### Cell culture, transfection, and C2C12 differentiation

HEK293T (293T), HeLa, and C2C12 cells were obtained from ATCC. All cells were cultured at 37 °C with 5% CO2 in DMEM supplemented with 10% FBS, 100 μg/mL streptomycin, and 100 U/mL penicillin. For glucose starvation, the cells were washed twice with PBS and then cultured in glucose-free DMEM supplemented with dialyzed FBS for 12-16 h, or in DM containing non-dialyzed 2% horse serum for 24 h. Cells were transfected with polyethylenimine (PEI) or Lipofectamine 3000 according to the manufacturer’s protocols. C2C12 myoblasts at 80%–90% confluency were induced to differentiate by switching GM to DM containing 2% horse serum. All cell lines were confirmed to be mycoplasma-free before the experiments.

### Antibodies, chemical reagents, and plasmids

The antibodies and reagents used in this study are listed in Supplementary Table S3. Full-length cDNA clones were prepared by RT_JPCR or purchased from the Bio-Research Innovation Center in Suzhou, BRICS, China. Truncations and point mutations were generated by standard molecular cloning methods. All plasmids used in this study are listed in Supplementary Table S3, and the oligos used for subcloning are listed in Supplementary Table S4.

### Animal work

C57BL/6J mice, Nono-KO mice, and Rosa26-CAG-LSL-Cas9-tdTomato mice (Cas9-KI) were purchased from Gempharmatech Co., Ltd., China. Mice were maintained in a pathogen-free environment. For Nono-KO mouse genotyping, tail genomic DNA was extracted using Buffer A (containing 25 mM NaOH and 0.2 mM EDTA) and Buffer B (containing 40 mM Tris-HCl, pH 8.0). Briefly, the samples were treated with Buffer A at 98 °C for 1 h, followed by the addition of Buffer B for PCR amplification. Nono-KO mice were identified by PCR amplification, yielding 362 bp and 461 bp products using specific primers targeting the NONO sequence, as listed in Supplementary Table S4. Cas9-KI mice were used for muscle-specific gene knockout. Adeno-associated virus 9 (AAV9) expressing Cre recombinase and sgRNA targeting Gys1 or Nono was administered by intramuscular injection. Specifically, 6- to 8-week-old male Cas9-KI mice received intramuscular (IM) injections of AAV9-sgRNAs at a dose of 5×10^11^ vector genomes (vg) per mouse. For CTX-induced muscle regeneration, the tibialis anterior (TA) muscle was injected with 50 μL of 20 µM CTX four weeks after AAV9 virus injection; control mice received PBS injections. TA muscles were harvested and analyzed at 0, 4, 7, 14, and 21 days post-CTX injection. The exercise tolerance test was performed as described previously [68]. The mice were acclimated (run for 9 min at 10 meters (m)/min followed by 1 min at 20 m/min at a 10° incline) on the treadmill for 2 consecutive days prior to the experimental protocol. Low-intensity (endurance) exercise studies were conducted as described previously [68]. In brief, fed mice were allowed to run for 10 min at 10 m/min, followed by a constant speed of 20 m/min at a 10° incline until exhaustion. Tail blood was collected after exercise, and lactate levels were measured using a Lactate Scout (Senelab, Germany) according to the manufacturer’s instructions. For mouse blood and tissue chemistry, blood glucose levels were determined using a OneTouch UltraMini glucose meter (OneTouch). Serum TG levels were determined using a triglyceride kit (Wako, 290-63701). Serum fatty acid levels were determined with a NEFA Kit (Wako, 294-63601). according to the manufacturer’s instructions. All animal experiments were performed following the ethical guidelines and protocols approved by the Institutional Animal Care and Use Committee (IACUC) of Sun Yat-sen University (Approve number: SYSU-IACUC-2022-002025).

### Immunoblotting (IB), immunoprecipitation (IP), and immunofluorescence (IF)

IB, IP, and IF experiments were performed as previously described [69]. For endogenous CoIP, 1 mg of cell lysate was precleared with protein A/G Sepharose (Beyotime Biotechnology, China) and then incubated with washed HA or Flag antibody-conjugated beads for 2 h overnight with rotation at 4 °C. The beads were collected and washed 3-5 times with lysis buffer, and 1 × loading buffer was added and boiled for 15 min at 95 °C to elute the bound proteins. For IF, cells were seeded on polylysine-coated glass coverslips and treated as indicated. The cells were then fixed with 4% paraformaldehyde (PFA) in PBS for 10 min and permeabilized with ice-cold methanol at 4 °C for 15 min or with 0.1% Triton X-100 at room temperature. After blocking with 5% bovine serum albumin (BSA), the cells were incubated with primary antibodies and subsequently with anti-rabbit or mouse secondary antibodies conjugated with Alexa Fluor 488/594/647. Nuclei were counterstained with DAPI (1 µg/mL) for 10 min, mounted with mounting media (90% glycerol in PBS), and stored at 4 °C until imaging.

### Glycogen isolation and content measurement

The cells were trypsinized and washed with cold PBS 3-5 times to remove residual glucose from the medium. For muscle or liver tissues, the samples were freshly homogenized, and protein lysate and glycogen were prepared from equal aliquots. For glycogen isolation, the sample was digested with 30% KOH for 2 h at 98 °C with occasional shaking. The glycogen was further precipitated in 66% ethanol for 24-48 h at -20 °C, after which the pelleted glycogen was washed and dissolved in ddH2O. Glycogen was quantified using an anthrone reagent kit (#BC0345, Solarbio Life Science, China) according to the manufacturer’s instructions.

### Histologic staining

For H.E. staining, the TA muscles were fixed in 4% paraformaldehyde, and the paraffin-embedded sections were stained according to standard procedures. The slides were covered with coverslips, mounted with xylene-based mounting media, and then scanned using a pathology slide scanner (KF-PR0-120-HI, KFBIO). For frozen section staining, freshly isolated tibialis anterior (TA) or gastrocnemius (GC) muscles were frozen in isopentane cooled in liquid nitrogen. 10 μm-thick serial muscle cross-sections were cut from the knee-cut side using a Leica CM1850 cryostat at -20 °C and mounted on positively charged glass slides.

Transverse sections collected from the widest part (mid-belly) of the GC muscle were used for histological comparison to maintain consistency between different mice. Frozen sections were stained with a Periodic Acid-Schiff (PAS) staining kit or with myogenic biomarker antibodies, following procedures similar to IF. WGA staining was performed using FITC-conjugated WGA. Notably, the gastrocnemius (GC) of each mouse was dissected as a whole. Quantification of the cross-sectional area of the myofibers was performed using ImageJ software, and histological staining quantification was conducted in a blinded manner with Image-Pro Plus software. For type I fiber staining quantification in the GC muscle, the data are presented as the number of type I fibers per medial head of the GC.

### Protein purification

Plasmids encoding Flag-AcGFP-GYS1, Flag-AcGFP-GYG1, Flag-mCherry-NONO, Flag-mCherry-FUS, or Flag-BFP-MyoD, were transfected into HEK 293T cells for protein purification. GYG1 (Y195F) was coexpressed with GYS1 by co-transfecting the GYS1 and GYG1 (Y195F) plasmids to enhance the solubility of GYS1. Cells were lysed in lysis buffer containing 50 mM Tris-HCl (pH 7.5), 1.0 M KCl, and 10% glycerol, supplemented with 1 mM dithiothreitol (DTT), protease inhibitors, and phosphatase inhibitors. The samples were sonicated and centrifuged at 12,000 rpm at 4 °C for 10 min. The supernatant was incubated with anti-Flag M2 affinity resin (A2220, Sigma) at 4 °C for 4 h with rotation, and the beads were subsequently washed four times with lysis buffer. Protein was eluted in lysis buffer containing 150 μg/mL 3xFlag peptide. The purified proteins were concentrated using Amicon® ultracentrifugal filters (Amicon Ultra 0.5, Merck) in storage buffer (50 mM Tris pH 7.5, 1.0 M KCl), flash-frozen in liquid nitrogen, and stored at -80 °C.

### *In vitro* droplet formation

*An in vitro* droplet formation assay was performed as previously described [70, 71]. Purified proteins were applied to 4-chamber glass-bottom microwell dishes (Cellvis, D35C4-20-1-N). Unused wells were filled with distilled water to maintain humidity in the nearby reaction chambers and prevent sample evaporation. 2–5 ul of purified proteins, or proteins mixed with RNA, glycogen, PEG800, or genomic DNA at the indicated concentrations, were gently dripped onto the glass surface of the microwell chambers, incubated for the indicated time, and photographed using a Nikon ECLIPSE Ti2 inverted microscope with a 60x oil objective.

### Image processing and statistical analysis

Images were processed using ImageJ (https://ImageJ.net/Fiji). Punctate cells were defined as those with more than five visible puncta in the nucleus or cytosol under the same exposure times. The quantified data are presented as the mean `±` SEM, and the statistical analysis methods are indicated in the figure legends. Statistical analyses were performed using GraphPad Prism 9. with p < 0.05 was considered to be statistically significant. All data from representative experiments were repeated at least three times independently, except those specifically noted in the figure legends. The uncropped western blot images are included in the supplementary file.

