## Supplementary material for "The metabolic enzyme GYS1 condenses with NONO/p54^nrb^ in the nucleus to spatiotemporally regulate glycogenesis and myogenic differentiation": Supplementary information.docx

**The supplementary material includes the following items:**

Supplementary methods

Supplementary Figures and Figure legends (9 Figures)

Supplementary references

Supplementary methods

Generation of stable cell lines, CRISPR/Cas9 gene knockout, and endogenous gene tagging

Stable cell lines were generated through lentiviral infection. 293T cells were transfected with pCDH constructs, pM2G, or psPAX2 packaging plasmids via the PEI method. 48 h post-transfection, the supernatant containing the virus was collected, passed through a 0.45 μm filter, and stored at −80 °C until use. For lentiviral transduction, cells were infected overnight with 8 μg/mL polybrene in virus-containing medium and selected with puromycin for 1-2 days post-infection. For CRISPR/Cas9 gene knockout, individual sgRNAs were subcloned and inserted into LentiCRISPRv2 at the BsmBI site, as described in the standard protocol, while dual sgRNAs were also subcloned and inserted into LentiCRISPRv2. Briefly, targeting cells were transfected with 2 µg of sgRNA plasmids for each 35-mm dish and selected with 1 µg/mL puromycin. 24-48 h after transfection, the puromycin-resistant cells were pooled, and single-cell clones were isolated by limiting dilution in a 96-well plate. For sequencing, genomic DNA was extracted from the clonal cells using lysis buffer (50 mM Tris, pH 8.0, 1 mM EDTA, 0.5% Tween 20, proteinase K > 0.6 U/mL), and amplification of the edited sequence from genomic DNA was performed using Taq PCR mix. The purified PCR products were sequenced to identify the correctly edited clones. For 3xHA endogenous tagging of GYS1 in 293T cells, the sgRNA and donor sequences were designed using the online tool TrueDesign Genome (https://apps.thermofisher.cn/apps/genome-editing-portal/#/summary). Targeting vector generation, gene targeting, and the identification of 3xHA-tagged GYS1 clones were performed as previously described [1]. The sgRNA sequences and primers used are listed in Supplementary Table S4.

Mass spectrometry analysis of the nGYS1 interactome by TurboID and TAP purification

TurboID proximal labeling was performed as previously described [2]. Briefly, HA-TurboID-GYS1-expressing 293T cells were labeled with 500 μM biotin at 37 °C for 10 min and then washed five times with ice-cold PBS. The cells were lysed in RIPA buffer supplemented with protease inhibitors and cleared by centrifugation. For biotinylated protein enrichment, 100 µL of washed PureProteome^TM^ streptavidin beads (LSKMAGT10, Millipore) were added to the clarified lysates, which were rotated for 1 h at room temperature and then incubated at 4 °C overnight. The beads were subsequently washed and sent for LC‒MS/MS analysis (MS Center for Excellence in Molecular Cell Sciences (Shanghai, China)). For immunoblotting and silver staining, 5% aliquots of beads were eluted by boiling in 50 µL of 3× SDS loading buffer supplemented with 20 mM DTT and 2 mM biotin. The resolved proteins were stained using a Pierce Silver Stain Kit (24612, Sigma) to verify the successful enrichment of the biotinylated material. For TAP purification, the cells were washed twice with PBS, and DSP (1 mM) was added to the cells and incubated at room temperature for 30 min with occasional agitation. Then, 0.2 M glycine in PBS was added to the cells for 15 min to terminate the cross-linking. After PBS washing, the cell nuclear fraction was extracted with high salt buffer (supplemented with protease inhibitors and 1 mM DTT). The nuclear extracts were then diluted with low salt buffer to 150 mM salt and subjected to Flag-IP with anti-Flag M2 beads (A2220, Millipore) for 2 h at 4 °C with rotation. The beads were washed four times with EBC buffer, and eluted with 150 μg/mL 3xFlag peptide in TBS, and the eluates were further subjected to Ni-NTA affinity purification with nickel beads (17-5248-05, GE Healthcare). The eluted samples were first silver-stained or immunoblotted to verify the purification and then shipped to Wininnovate Bio. Company (Shenzhen, China) on ice for LC‒MS/MS analysis. For GO pathway analysis of the identified interacting proteins, the R package “clusterProfiler” was used.

Cytosolic and nuclear protein fractionation

Cells were first lysed with low salt buffer (10 mM HEPES; pH 7.5–8.0, 10 mM KCl, 0.1 mM EDTA, 1 mM DTT, supplemented with protease and phosphatase inhibitors) for 30 min at 4 °C. Then, 100 U/mL amylase was added to the sample with constant rotation at room temperature for 15 min to release GYS1 from dissolvable glycogen. Nuclei were pelleted by centrifugation at 1000 × g for 10 min at 4 °C, and the supernatant containing cytoplasmic proteins (Cyto) was collected. The remaining nuclei were washed three times with low salt buffer, lysed in high salt buffer (20 mM HEPES; pH 7.5–8.0, 400 mM NaCl, 1 mM EDTA, 1.5 mM MgCl2, 0.1% SDS, and 1 mM DTT, supplemented with protease and phosphatase inhibitors) on ice for 30 min and sonicated for 3x 10 seconds at 20% output. Supernatants containing soluble nuclear proteins (Nucl.) were collected by centrifugation at 20, 000 × g for 10 min and stored at -80 °C before analysis.

RT‒qPCR, RIP‒qPCR, and ChIP‒qPCR

Total RNA was extracted using TRIzol according to a standard RNA extraction protocol. Reverse transcription was performed with 1 μg of total RNA using a 5X Evo M-MLV RT Reaction Mix Ver.2 kit (AG11728, Accurate Biology) according to the manufacturer’s instructions. RT‒qPCR analyses were performed using StepOnePlus with 2× RealStar Fast SYBR qPCR Mix (A303, GenStar). mRNA levels were quantified using the ΔΔCt method, and the results were normalized to those of β-actin or GAPDH. RIP-qPCR was performed as previously described [3]. Briefly, 3×HA endogenous tagged GYS1 293T cells were glucose-starved and lysed in polysome lysis buffer (100 mM KCl, 5 mM MgCl2, 10 mM HEPES (pH 7.0), 0.5% NP40, 1 mM DTT) supplemented with 100 U/mL RNase Out (10777019, Invitrogen), protease inhibitors, and 200 μM VRC. The cleared lysates were then subjected to IP with HA beads (P2121, Beyotime Biotechnology). The specifically enriched RNA components were released by adding proteinase K to the beads and extracted using the standard TRIzol extraction protocol.

Flag-AcGFP-MyoD ChIP was performed as previously described [4]. Briefly, approximately 1×10^7^ C2C12 cells were cross-linked with 1% formaldehyde (10 min, RTM) and quenched with glycine. The cells were subsequently washed and harvested by centrifugation (700 g, 5 min at 4 °C). Afterward, the cells were lysed in 1 mL of lysis buffer 1 (50 mM HEPES-KOH pH 7.5, 150 mM NaCl, 1 mM EDTA, 0.25% Triton X-100, 0.5% NP-40, 10% glycerol, supplemented with complete protease inhibitors) and incubated on a rotator at 4 °C for 10 min. Following centrifugation at 1350 g for 5 min at 4 °C, the pellets were washed with 10 mL of lysis buffer 2 (10 mM Tris-HCl pH 8.0, 200 mM NaCl, 1 mM EDTA pH 8.0, 1 mM EGTA pH 8.0, supplemented with complete proteinase inhibitors) by incubation on a rotator at 4 °C for 10 min. The nuclei were rinsed twice with 2 mL of sonication buffer (10 mM Tris-HCl pH 8.0, 0.1% SDS, 1 mM EDTA, supplemented with complete proteinase inhibitors) and sonicated with a Covaris S220 (intensity 140 W, duty cycle 5%, cycles per burst 200, 8 min). The resulting lysate was then supplemented with NaCl and Triton X-100 to achieve final concentrations of 150 mM NaCl and 1% Triton X-100 and cleared by centrifugation for 30 min at 17,000 g. Finally, the lysate was incubated with washed Dynabeads Protein A (AS046, ABclonal) for 2 h at 4 °C. The cleared lysate was then incubated with 3 μg of anti-Flag antibody (SA042005, Smart Life Sciences) or IgG control (sc-2027, Santa Cruz) overnight at 4 °C. The following day, 30 μL of washed Dynabeads Protein A was added to each immunoprecipitation (IP), followed by further incubation at 4 °C for 3 h. Subsequently, the beads were collected and subjected to the following washing steps: two washes with IP buffer, two washes with high-salt wash buffer (containing 10 mM Tris-HCl pH 8.0, 500 mM NaCl, 0.1% SDS, 1 mM EDTA, 1% Triton X100), two washes with LiCl wash buffer (containing 10 mM Tris-HCl pH 8.0, 250 mM LiCl, 1 mM EDTA, 0.5% NP-40, 0.5% deoxycholate), and finally, one wash with cold TE buffer (containing 10 mM Tris-HCl pH 8.0, 1 mM EDTA, 50 mM NaCl). The immunocomplexes were eluted in ChIP elution buffer (composed of 50 mM Tris-HCl pH 8.0, 10 mM EDTA, 1% SDS) at 65 °C for 30 min, followed by reverse cross-linking through incubation at 65 °C for 16 h. Immunoprecipitated DNA was then treated with RNase A (0.2 mg/mL) at 37 °C for 2 h, followed by proteinase K treatment (0.2 mg/mL) at 55 °C for 2 h. Subsequently, the DNA was subjected to phenol:chloroform:isoamyl alcohol (P:C:IA) extraction and ethanol precipitation. The extracted DNA served as a template for qPCR using multiple pairs of primers targeting the genomic locus of interest, and the percentage of input recovery was calculated accordingly. All experiments were performed in triplicate. The PCR primers used are listed in Supplementary Table S4.

AAV9 construct modification, virus packaging, and purification

The pAAV-TBG-*cre-Luci*-sgRNA vector was kindly provided by Prof. Qiurong Ding (Shanghai Institute of Nutrition and Health). The plasmid was modified by replacing the original TBG promoter with the MH promoter (synthesized by Tsingke, China) for muscle cell-specific expression, and the original T2A-luciferase-sgRNA cassette was replaced with the P2A-Nluc-sgRNA cassette (synthesized by Tsingke, China). The modified plasmid pAAV-MH-*Cre-Nluc*-sgRNA was used to subclone two sgRNA sequences targeting *Nono* or *Gys1*, or as a control. AAV9 viruses were generated by transfecting the packaging plasmids pAAV helper, pAAV9 package plasmid, and pAAV-MH-*Cre-Nluc*-sgRNA (Vector, *Gys1-*sgRNA1*-*sgRNA2 or *Nono-*sgRNA1-sgRNA2) into 293T cells using PEI. For AAV9 purification, the virus supernatant was collected by centrifugation at 4000 rpm for 20 min at 4 °C and concentrated to 10-15 mL using a concentration column (Merck UFC905096). Then, ioxanol (Sigma‒Aldrich, Optiprep, D1556) gradient concentrations were prepared by layering 3.5 mL of 60%, 3.5 mL of 40%, 4 mL of 25%, and 4 mL of 17% ioxanol into centrifuge tubes from bottom to top. The concentrated virus supernatant and lysed cell supernatant were then gently added to the top layer of the centrifuge tube and centrifuged at 60,000 rpm for 3 h at 4 °C using a 70Ti rotor (Beckman OPTIMAL-80XP). The layer containing the virus at a concentration of 40% was transferred to a new tube for concentration, followed by three washes with 10 mL of PBS at 4000 rpm for 20 min at 4 °C. The virus was then concentrated to a final volume of 500 µL.

Live-cell microscopic imaging and fluorescence recovery after photobleaching (FRAP)

The cells were grown in 4-chamber glass-bottom microwell dishes (Cellvis, D35C4-20-1-N). After transfection with fluorescent protein-tagged plasmids as indicated, live-cell imaging was performed using a Nikon ECLIPSE Ti2 inverted microscope with a 60x oil objective. For time-lapse imaging, the cells were placed in an incubation chamber maintained at 37 °C with 5% CO2. The exposure time was approximately 100 ms, and images were taken at regular intervals. Confocal images were obtained using a Zeiss LSM 880 confocal microscope with a 60x oil objective. For the FRAP assay, cells were plated on 35-mm glass bottom dishes and transfected with GFP- or mCherry-fusion plasmids for 24–36 h before imaging. FRAP was performed with a Zeiss LSM 880 confocal microscope with a 60x oil immersion objective (Plan-Apochromat, 1.4 Oil DIC M27). The cells were maintained at 37 °C and 5% CO2 in a humidified stage-top incubator during the experiment. Bleaching was conducted with a pulse 488 nm laser or a 561 nm laser at 70%–80% laser power, and time-lapse images were acquired at regular intervals after bleaching.

Stochastic optical reconstruction microscopy (STORM)

HeLa cells or HEK 293T cells expressing Flag-GYS1 and HA-NONO were seeded on glass-bottom dishes for IF staining. The imaging buffer was prepared according to the protocol provided by Nikon. Briefly, glucose oxidase (G2133, Sigma) and catalase (C1345, Sigma) stock solutions were dissolved in Buffer A (10 mM Tris, 50 mM sodium chloride, pH 8.0) at final concentrations of 70 mg/mL and 17 mg/mL, respectively. GLOX solution was freshly prepared by mixing glucose oxidase and catalase stock solutions at a 4:1 ratio. The imaging buffer was freshly prepared by mixing 7 μL of GLOX, 7 μL of 2-mercaptoethanol (07604, Sigma), and 690 μL of Buffer B (50 mM Tris, 10 mM sodium chloride, 10% glucose, pH 8.0), which was added to the glass-bottom dish before imaging. Fluorescence signals of Alexa Fluor 568-labeled anti-Flag or anti-GYS1 and Alexa Fluor 647-labeled anti-HA or anti-NONO were detected sequentially on a Nikon N-STORM microscope system using a 100× oil objective lens (APO TIRF, 1.49 NA, Nikon) and an ORCA-Flash 4.0 V2 sCMOS camera (Hamamatsu) with 40 ms exposure in the non-binning mode. Flag-GYS1 or GYS1 was excited with a 561 nm laser, and fluorescence was detected in the RFP channel. The photoconversion of Alexa Fluor 568 was triggered by a 405 nm laser, with the intensity gradually increasing until all Alexa Fluor 568 signals were photoconverted and detected. For imaging HA-NONO or endogenous NONO, Alexa Fluor 647 signals were excited with a 640 nm laser, and fluorescence was detected in the Cy5 channel. The “On”-“off” photoswitching of Alexa Fluor 647 occurred spontaneously, with a 405 nm laser occasionally used to accelerate the photoswitching. Single-molecule localization and image reconstruction were performed using Nikon NIS Element software (v5.1) with the N-STORM plugin.

Supplementary figures and figure legends

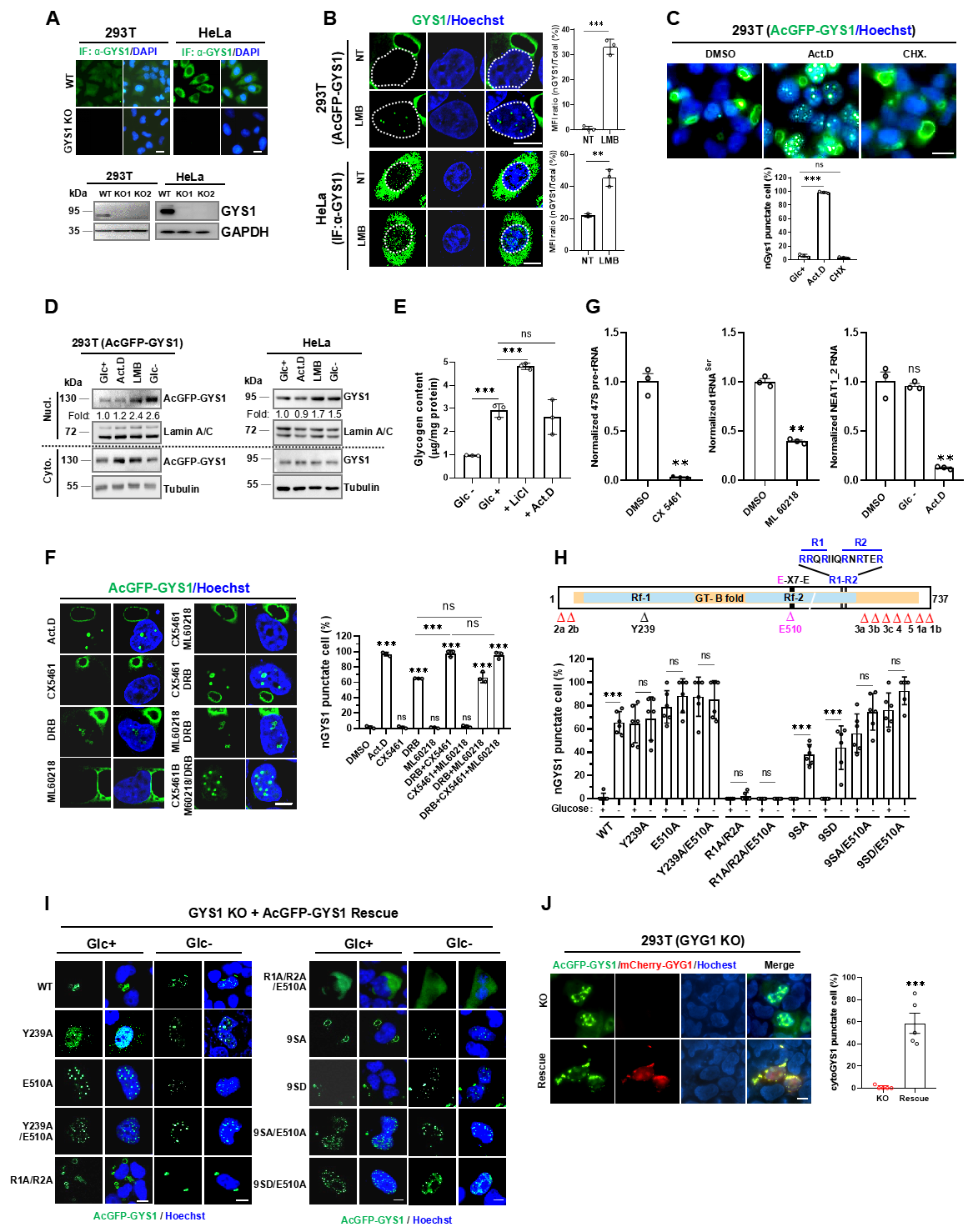

Fig. S1. Regulation of GYS1 subcellular compartmentalization. A Validation of the GYS1 antibody by IF and IB in 293T and HeLa GYS1-KO cells generated by CRISPR/Cas9 gene editing. B Imaging and quantification of subcellular GYS1 protein bodies after LMB treatment (100 nM, 12 h). AcGFP-GYS1 live cells or GYS1 IF staining after the indicated treatment in 293T or HeLa cells (left), and the mean fluorescence intensity (MFI) of nGYS1 was quantified by the MFI ratio between the nGYS1 and total GYS1 signals (right). C Imaging and quantification of nGYS1 protein bodies with Act.D or CHX (20 μg/mL) treatment for 12 h. D IB analysis of the subcellular GYS1 distribution by cell fractionation. AcGFP-GYS1-expressing 293T cells or HeLa cells with indicated treatments were fractionated into cytosolic (cyto.) and nuclear (Nucl.), and IB analyzed with indicated antibodies. E Glycogen content in 293T cells treated with glucose starvation, 20 mM LiCl, or 1 μg/mL Act.D for 12 h. F Imaging and quantification of nGYS1 puncta with RNA Pol inhibitors in 293T cells. Live cell imaging of AcGFP-GYS1 cells with indicated treatment (Act.D, 1 μg/mL; CX5461, 10 μM; ML60218, 20 μM) (left); quantification of nGYS1 punctate cells (>5 puncta per nucleus) of each treatment (right). G RT-qPCR analysis of RNA levels post-treatment with RNA Pol inhibitors or glucose starvation. **H** A diagram showing GYS1 domain structure and key functional sites (top) and quantified nGYS1 punctate cells (bottom). I Representative images of AcGFP-GYS1 mutants in GYS1-KO cells. **J** Imaging of subcellular GYS1 distribution in GYG1-KO and mCherry-GYG1-rescued cells (left). The percentage of cytosolic GYS1 punctate cells was quantified in five random fields of view (FOVs) (right). Error bars show the mean ± SEM; statistical analysis using two-tailed t-test; scale bar, 10 μm.

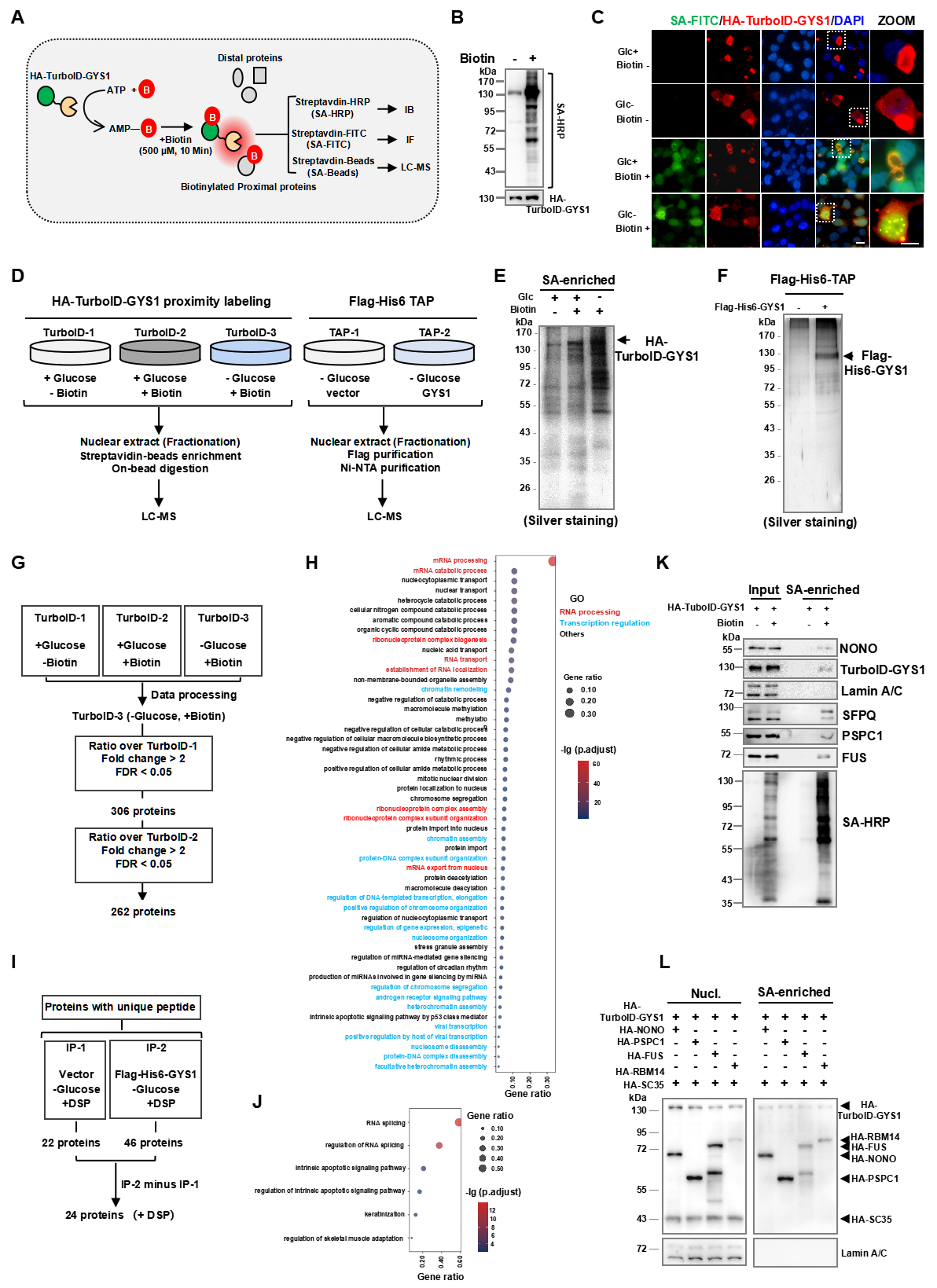

Fig. S2. Identification and validation of GYS1-interacting proteins in the nucleus. A Schematic diagram of HA-TurboID-GYS1 proximity labeling for method validation and MS identification of the nGYS1 interactome. B IB validation of biotinylated proteins with the SA-HRP antibody in 293T cell lysates. C IF staining of biotinylated proteins in the nuclei of normal cultured or glucose-starved 293T cells. Biotinylated proteins and HA-TurboID-GYS1 were stained with SA-FITC and HA antibodies, respectively. D Workflows for the identification of nGYS1-interacting proteins by turboID-MS and TAP-MS in 293T cells. E Silver staining of SA-bead-enriched proteins labeled by TurboID-GYS1 for MS analysis. F Silver staining of purified proteins by Flag and His two-step TAP. G Data processing workflow for the identification of nuclear GYS1 proximal proteins. H Gene ontology pathway analysis of nGYS1-interacting proteins identified by TurboID-MS. I Data processing workflow for identifying nuclear GYS1-interacting proteins by TAP-MS. J The enriched gene ontology pathways for nGYS1-interacting proteins identified by TAP-MS. K Proximity labeling of paraspeckle protein by TurboID-GYS1 in 293T cells. Biotinylated proteins were enriched by SA beads from glucose-starved TurboID-GYS1-expressing 293T cell lysates and analyzed by IB with the indicated antibodies. L Proximity labeling of paraspeckle proteins by TurboID-GYS1 in 293T cells. Cells were transfected with various paraspeckle protein cDNAs as indicated; SC35 was used as an internal negative control, and the SA-bead enriched biotinylated proteins were analyzed with the indicated antibodies; scale bar, 10 μm.

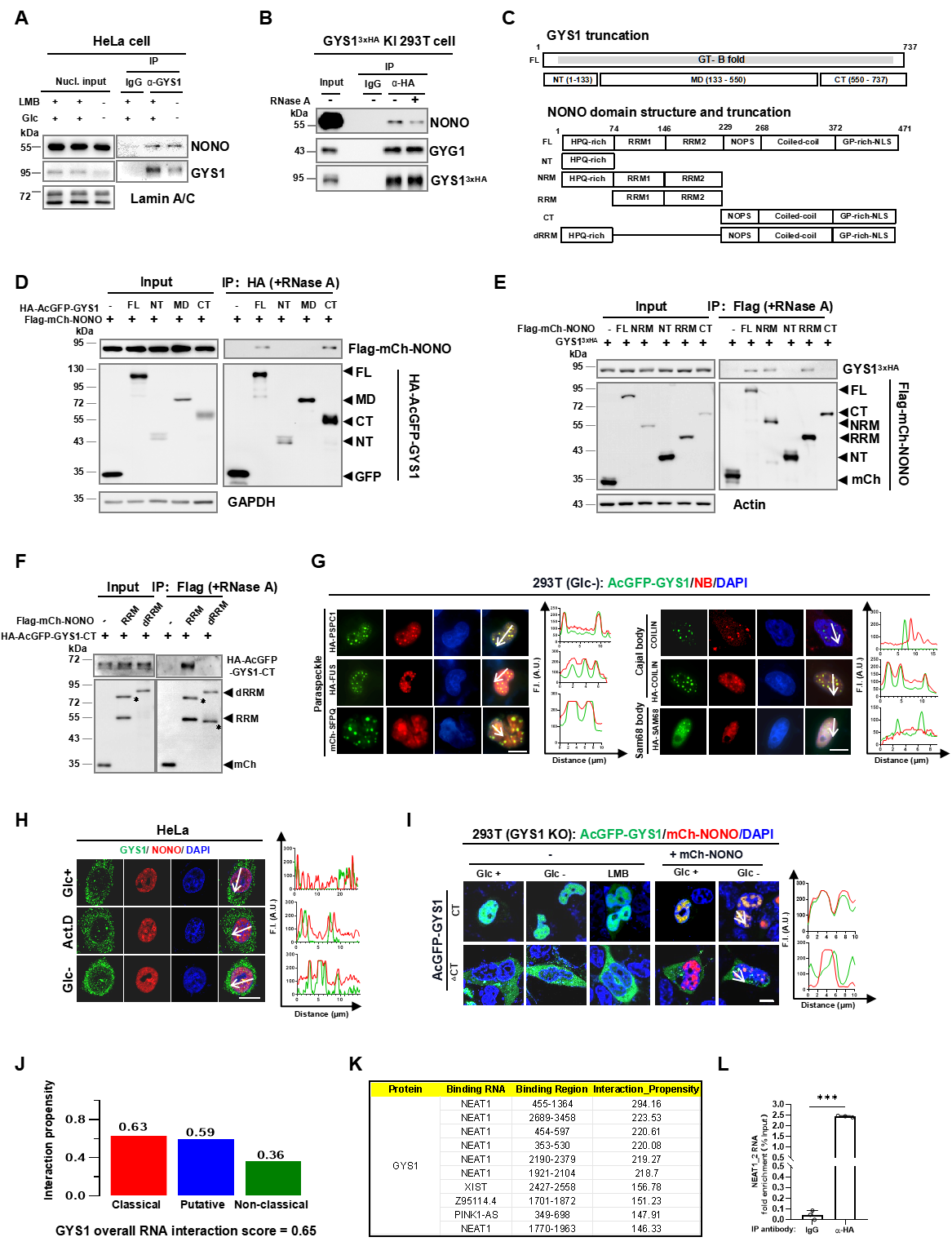

Fig. S3. nGYS1 is an RBP that interacts and compartmentalizes with NONO. A Co-IP validation of the NONO-GYS1 interaction in HeLa cells by GYS1 IP, 100 nM LMB, or glucose starvation for 12 h. B Co-IP analysis of the NONO-GYS1 interaction in GYS1^3xHA^ 293T KI cells treated with RNase A (100 μg/mL, 4 °C, 1 h). C Truncation and domain structures of GYS1 and NONO. D Co-IP analysis of the interaction between NONO and GYS1 truncation mutants rescued in GYS1-KO 293T cells, and the HA-IPs were treated with RNase A. E Co-IP analysis of the interaction between GYS1^3xHA^ and various NONO truncations in Nono-KO 293T cells, and the Flag-IPs were treated with RNase A. F Co-IP analysis of the interaction between GYS1-CT and NONO-RRM in NONO/GYS1-DKO 293T cells, and the Flag-IPs were treated with RNase A. G Colocalization of nGYS1 and nuclear body proteins. AcGFP-GYS1 was coexpressed with HA-tagged NB markers and IF-stained after 12 h glucose starvation. Colocalization was analyzed by the fluorescence intensity profile along a line crossing the nucleus. H Endogenous GYS1 and NONO colocalization in HeLa cells with the indicated treatments. I Subcellular localization of AcGFP-GYS1-CT or △CT in response to Glc-, LMB, and their colocalization with mCh-NONO in 293T cells. J RNA-binding propensity of GYS1 predicted by catRAPID. K The top ten GYS1-binding lncRNAs predicted by catRAPID. L Validation of the GYS1-NEAT1_2 interaction by RIP-qPCR in glucose-starved GYS1^3xHA^ 293T-KI cells. Native RIP was performed with IgG or HA antibody, and RT-qPCR was performed to analyze the enrichment of NEAT1_2. Error bars show the mean ± SEM; statistical analysis was performed using two-tailed t-test; scale bar, 10 μm.

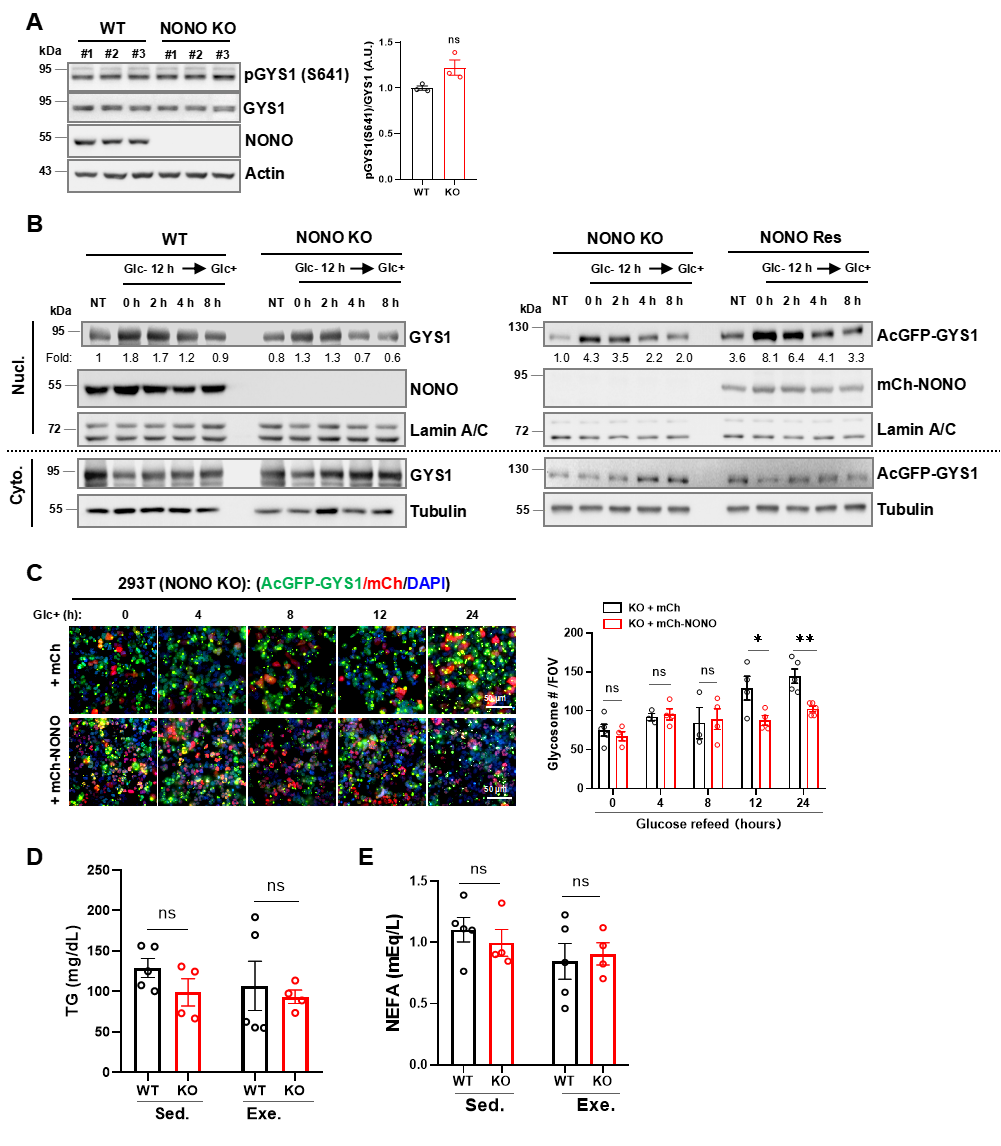

Fig. S4. NONO retains GYS1 in the nucleus to inhibit glycosome formation. A IB analysis of GYS1 phosphorylation in WT and NONO-KO 293T cells. B IB analysis of nGYS1 export upon glucose restimulation in WT, NONO-KO, and NONO-rescued 293T cells. NT, normal culture. C Glycosome GYS1 in NONO-KO or NONO-rescued cells. Imaging of vector- or mCherry-NONO-rescued AcGFP-GYS1-expressing NONO-KO 293T cells with 25 mM glucose restimulated for the indicated time after 12 h of glucose starvation (left). AcGFP-GYS1-labeled glycosomes were quantified in three FOVs at each time point (right); D, E Blood TG (D) and NEFA (E) levels in the Sed. or Exe. mice. n = 4-5 mice per group. Error bars show the mean ± SEM; statistical analysis was performed using two-tailed t-test; scale bar, 50 μm.

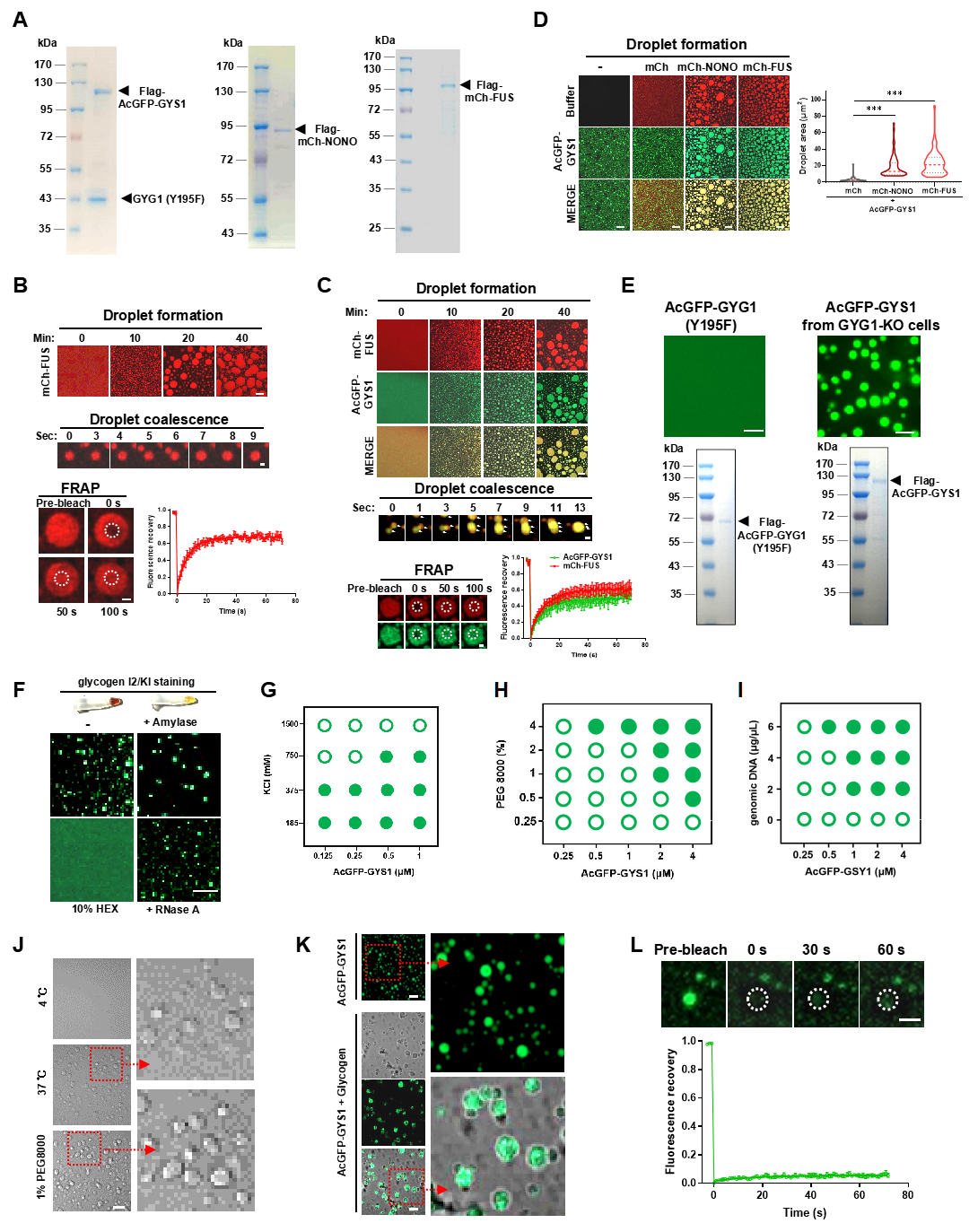

Fig. S5. GYS1 protein purification and phase separation with RBP partners *in vitro.* A Coomassie blue staining of the purified Flag-AcGFP-GYS1/GYG1(Y195F) complex (left), Flag-mCherry-NONO (middle), and Flag-mCherry-FUS (right). B LLPS of 3 μM purified mCherry-FUS (top) and droplet coalescence (middle). FRAP assay of FUS (bottom left); scale bars, 2 µm. The FRAP signals were quantified from n = 6 droplets (bottom right). The buffer conditions were 50 mM Tris-HCl (pH 7.5), 150 mM NaCl at 25 °C. C Co-condensation of mCherry-FUS (3 μM) and AcGFP-GYS1 (4 μM) mixtures (top) and droplet coalescence (middle); scale bars, 2 µm. FRAP assay of FUS and GYS1 mixture droplets (bottom left); scale bars, 2 µm. The FRAP signals were quantified from n = 7 droplets (bottom right). D The purified mCherry or mCherry fusion protein was mixed with AcGFP-GYS1 to a final concentration of 3 μM for each protein. After 10 min, the droplet size in four random FOVs, about 60 droplets for each FOV, was measured, and the size was quantified using violin plots generated by the particle analysis algorithm of imageJ; red line, median; black line, interquartile range; statistical analysis was performed using unpaired t-test. E Droplet formation of 4 μM purified Flag-AcGFP-GYG1(Y195F) or 4 μM Flag-AcGFP-GYS1 purified from transfected GYG1-KO cells. F AcGFP-GYS1 droplet formation with buffer (-), HEX (10%), RNase A (0.1 μg/μL), and amylase (10 units/μL), and the reactions were incubated for 30 min before imaging. The tubes show I2/KI staining of 1 μg/μL glycogen with or without 1 unit/ul amylase treatment. G-I Phase diagrams of AcGFP-GYS1 with varying concentrations of protein and KCl (G), PEG8000 (H), or purified genomic DNA (I); filled circle, droplet formation; empty circle, no droplet formation. J Bright-field images of glycogen droplets. Assays were performed with 5.0 μg/μL freshly purified mouse liver glycogen, and the reactions were incubated for 30 min under the indicated conditions in buffer containing 50 mM Tris-HCl (pH 7.5), 150 mM NaCl. K Droplet formation of AcGFP-GYS1 (4 μM) alone or glycogen-AcGFP-GYS1 mixture (5 μg/μL glycogen with 4 μM GYS1). The droplets were visualized in bright fields and 448 nM channels 30 min after mixing. Assay buffer: 50 mM Tris-HCl (pH 7.5), 150 mM NaCl, and 1% PEG8000. L FRAP assay of glycogen-AcGFP-GYS1 droplets (top) and quantification of 5 droplets (bottom). Error bars represent the mean ± SEM; scale bar, 10 μm unless otherwise specified in each image.

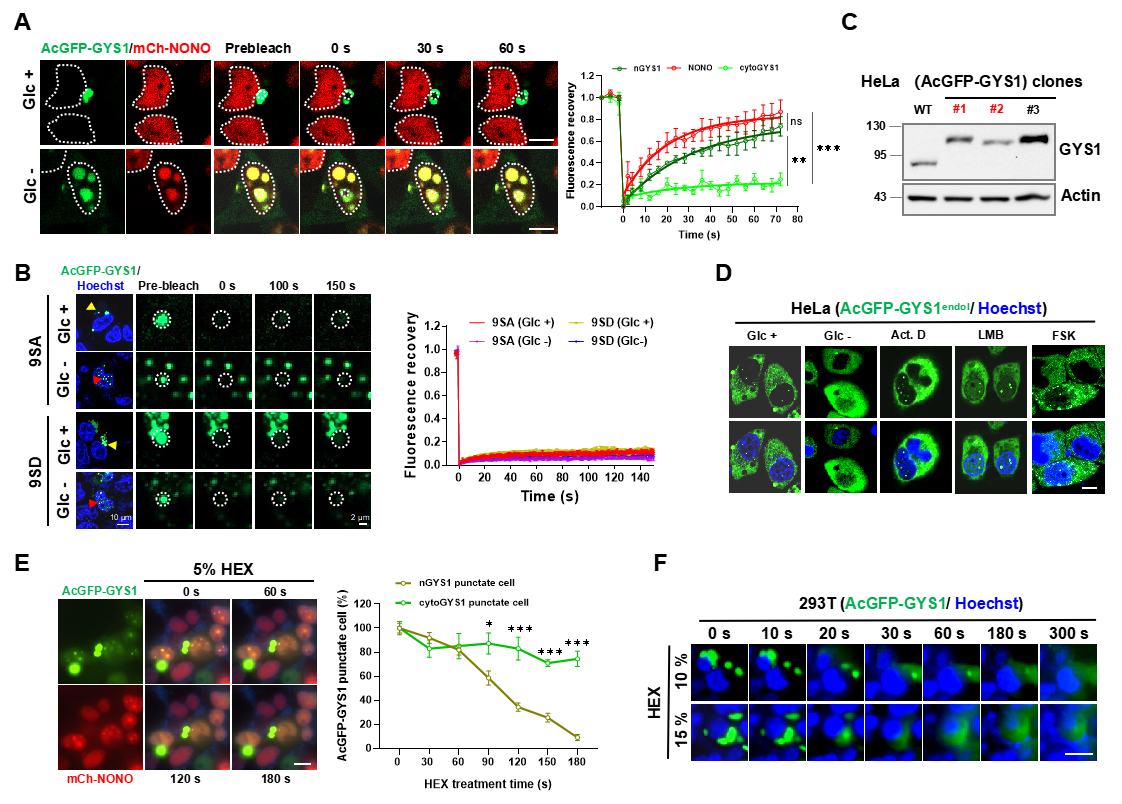

Fig. S6. Different dynamics of GYS1 protein bodies in the nucleus and cytosol. A FRAP assay of the subcellular distinct microbodies of AcGFP-GYS1 and mCherry-NONO with glucose starvation (left) and quantified FRAP signals for each type of puncta (n= 5) (right). The data were analyzed using one-way ANOVA with multiple comparisons. B Representative images and FRAP assay for nuclear and cytosolic puncta of 9SA or 9SD mutants rescued in 293T GYS1-KO cells treated with glucose starvation (left) and quantified FRAP signals from 5-8 puncta of each type (right). C IB analysis of AcGFP-GYS1 expression in HeLa clonal cells. The clones numbered in red denote the endogenous-level expression of the protein. D Imaging AcGFP-GYS1^endol^ nuclear puncta formation after various treatments (1 μg/mL Act.D, 100 nM LMB, 100 μM FSK, or glucose starvation for 12 h). E Diffusion of subcellular distinct microbodies of AcGFP-GYS1 with or without mCherry-NONO by 5% HEX treatment at the indicated time points (left) and quantified punctate cells of each type, n = 3-4 FOVs, with about 50 cells in each FOV (right); nGYS1 punctate cells and cytoGYS1 punctate cells were quantified and compared using multiple t-test (right). F Diffusion of cytosolic AcGFP-GYS1 puncta after 10% or 15% HEX treatment at the indicated times. Error bars show the mean ± SEM; scale bar, 10 μm unless otherwise specified in each image.

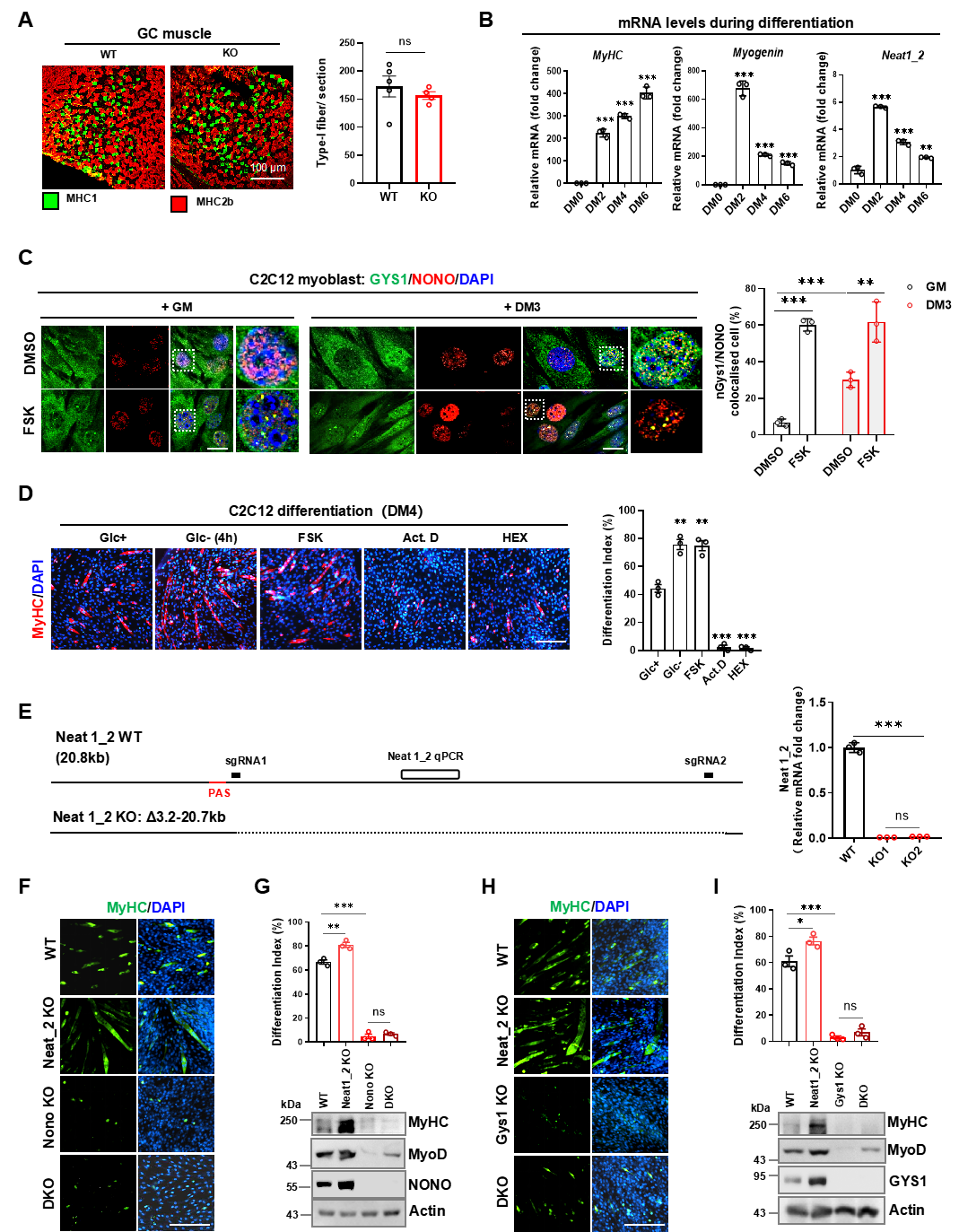

Fig. S7. Gys1 and Nono are pivotal for C2C12 myoblast differentiation and myofiber development. A Representative images and quantification of MHC1 (green) and MHC2b (red) staining of GC muscles from 8-week-old male mice of the indicated genotypes. n = 4–5 mice per group. B RT-qPCR analysis of the mRNA levels of the myogenic genes *MyHC*, *Myogenin*, and *Neat1_2* during C2C12 differentiation. C Gys1-Nono colocalization in C2C12 cells cultured in GM or DM for 3 days (DM3) and the response to FSK (100 μM, 12 h) (left). Colocalized cells were quantified by counting the percentage of cells with more than five colocalized puncta in the nucleus; three FOVs were analyzed ed for each treatment. D Effects of various pretreatments on C2C12 differentiation. Before switching to DM culture, cells were pretreated for 4 h with glucose starvation, FSK, Act.D, and 0.5% HEX. **E** Schematic depicting the *Neat1_2* gene structure, the design of the sgRNA, and the indicated qPCR amplicon (left); RT-qPCR analysis of *Neat1_2* mRNA levels in WT and *Neat1_2-*KO C2C12 cells (right). F-I Differentiation of Neat1_2-KO or DKO C2C12 cells. MyHC staining of WT, Neat1_2-KO, NONO-KO, and Neat1_2/Nono double-KO cells (DKO) (F) or GYS1-KO and Neat1_2/Gys1 DKO cells (H) 4 days after differentiation. The differentiation index was calculated by the ratio of the number of nuclei in the myotubes to the total number of nuclei in one FOV (G, I top), and MyHC protein levels were IB analyzed in C2C12 myoblasts of the indicated genotypes (G, I bottom). Error bars show the mean ± SEM; statistical analysis using unpaired t-test; scale bars: 100 μm (A), 10 μm (C), and 50 μm (E, F, H).

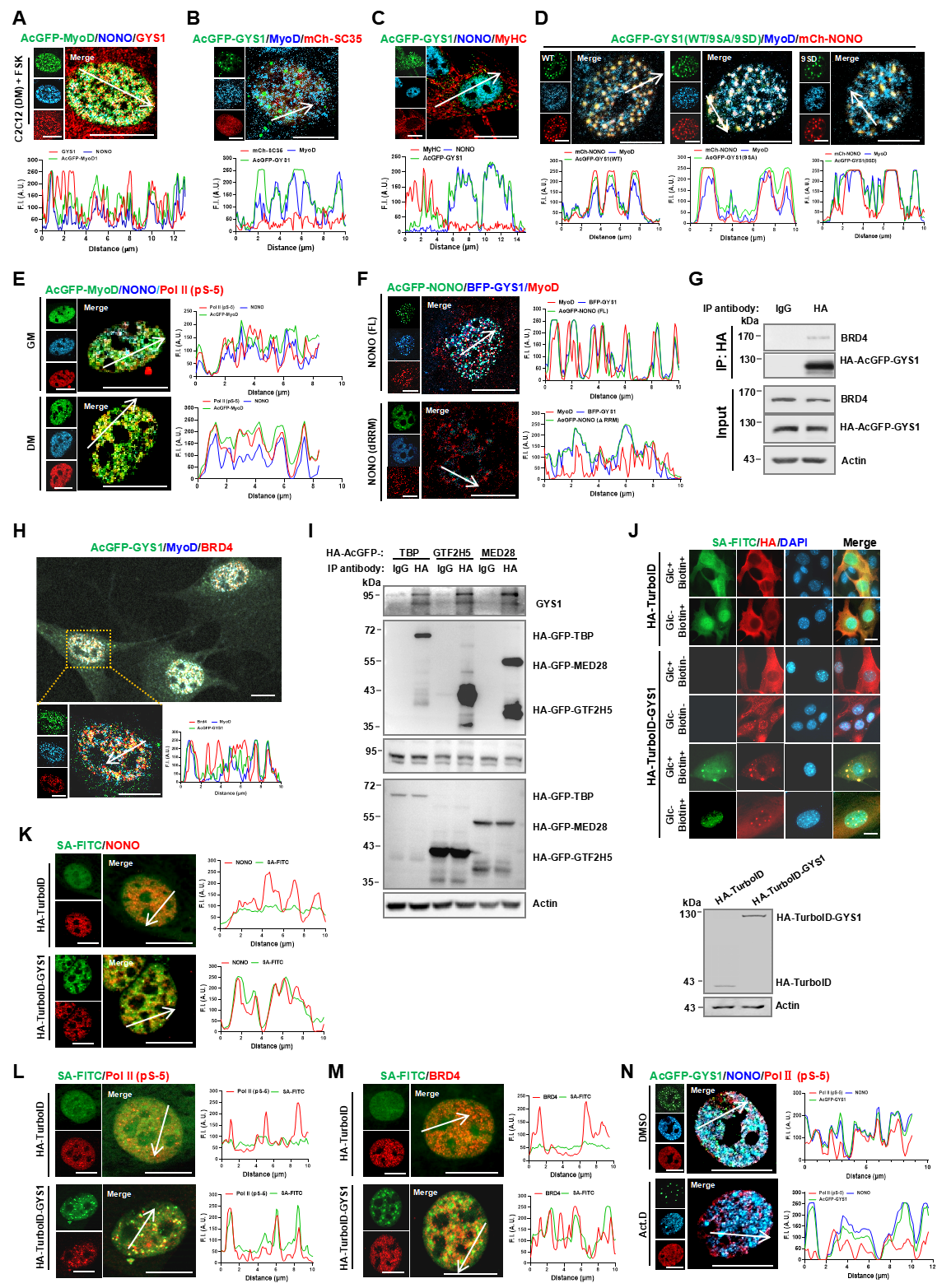

Fig. S8. GYS1-NONO co-condenses MyoD and PIC into transcriptional condensates during C2C12 differentiation. A Colocalization of MyoD, Nono, and Gys1 in C2C12 cells treated with FSK (100 μM, 1 h). B Colocalization of AcGFP-GYS1, mCherry-SC35, and MyoD. **C** Colocalization of AcGFP-GYS1, Nono, and MyHC. **D** Colocalization of AcGFP-GYS1 WT, 9SA, and 9SD with MyoD and mCherry-NONO. E Colocalization of AcGFP-MyoD, Nono, and Pol II (pS-5) in C2C12 cells cultured in GM or DM for 24 h. **F** Colocalization of AcGFP-NONO full length (FL) or dRRM with MyoD and BFP-GYS1 in Nono-KO C2C12 cells. **G** Co-IP analysis of GYS1 interactions with Brd4 in C2C12 cells. **H** Colocalization of AcGFP-GYS1 with MyoD and Brd4. **I** Co-IP analysis of GYS1 interactions with TBP, GTF2H5, and MED28 in C2C12 cells. **J** Verification of TuroID-GYS1 proximity labeling by IF staining of biotinylated proteins with SA-FITC in the nuclei of C2C12 cell lines treated as indicated. The stably expressed HA-tagged TurboID or TurboID-GYS1 was verified by WB. **K-M** TuroID-GYS1 proximity labeling of PIC through IF staining of biotinylated NONO (**K**), active Pol II (pS-5) (**L**), and Brd4 (**M**) with SA-FITC and specific antibodies in the nuclei of C2C12 cell lines. **N** Colocalization of AcGFP-GYS1, Nono, and Pol II (pS-5) with or without Act.D treatment (1.0 μg/mL, 4 h). Cells were treated with glucose-free DM for 24 h to deplete glucose from horse serum before staining or lysis unless specified. Scale bars: 10 μm.

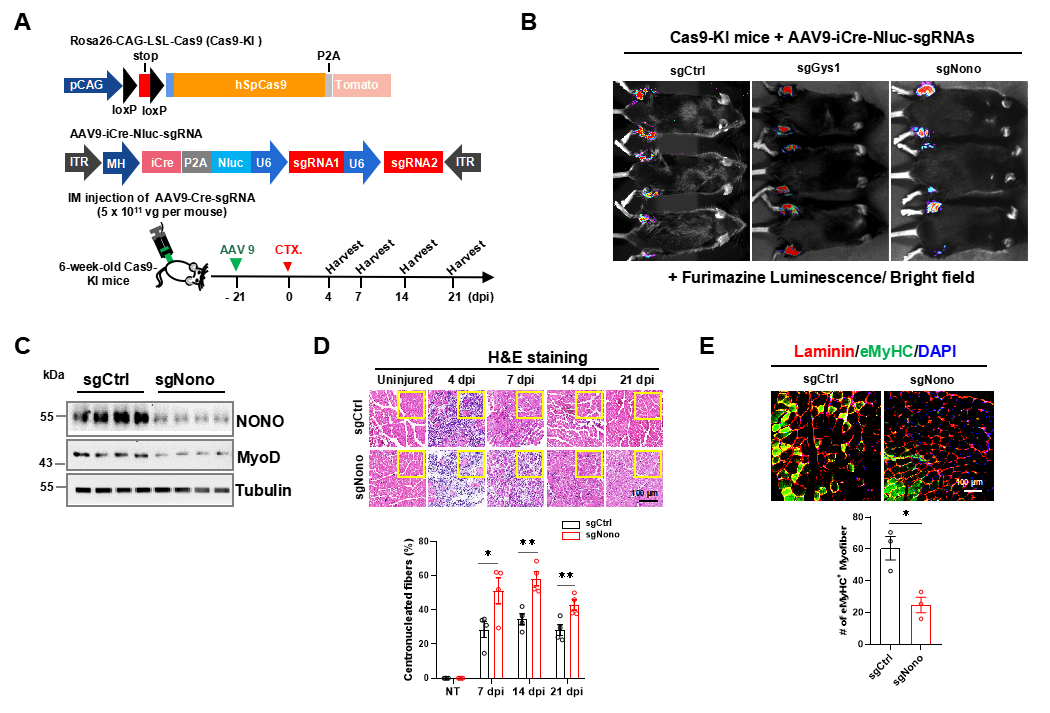

Fig. S9. nGys1-Nono regulates CTX-induced muscle regeneration in mice. A Diagram of the Cas9 KI locus (top), the AAV9 construct for Cre-Nluc-sgRNA (middle), and a schematic outlining the experimental timeline for CTX-induced muscle regeneration (bottom). B *In vivo* bioluminescence imaging of live mice injected with the indicated Cre-expressing virus. C IB analysis of Nono knockdown muscle tissues 28 days after AAV9 injection. D H&E staining of TA muscle cross-sections from AAV9-infected muscles at 0, 4, 7, 14, and 21 dpi (top), and quantified centronucleated myofibers at the indicated time points (bottom); myofibers containing centralized nuclei were quantified from 200 myofibers of the TA muscle in each mouse using ImageJ. E Laminin and eMyHC immunostaining of TA muscles at 7 dpi (top) and the percentage of eMyHC^+^ myofibers in 3 FOVs/section (bottom); n = 5-6 mice per group. Error bars show the mean ± SEM; statistical analysis using unpaired t-test; scale bars: 100 μm (D, E).
