## Supplementary figures and images for "The metabolic enzyme GYS1 condenses with NONO/p54^nrb^ in the nucleus to spatiotemporally regulate glycogenesis and myogenic differentiation"

### Fig. S1.tif

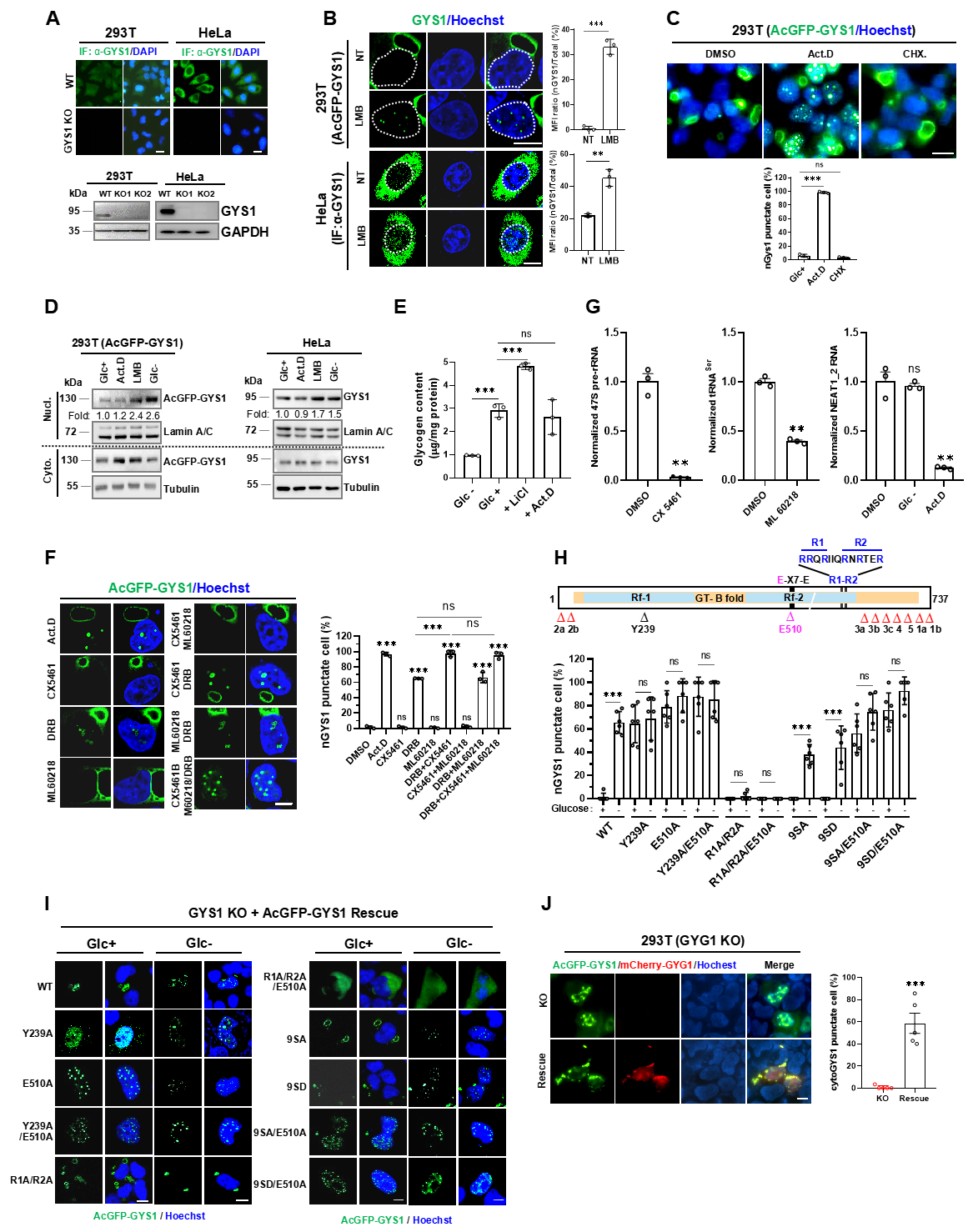

### Fig. S2.tif

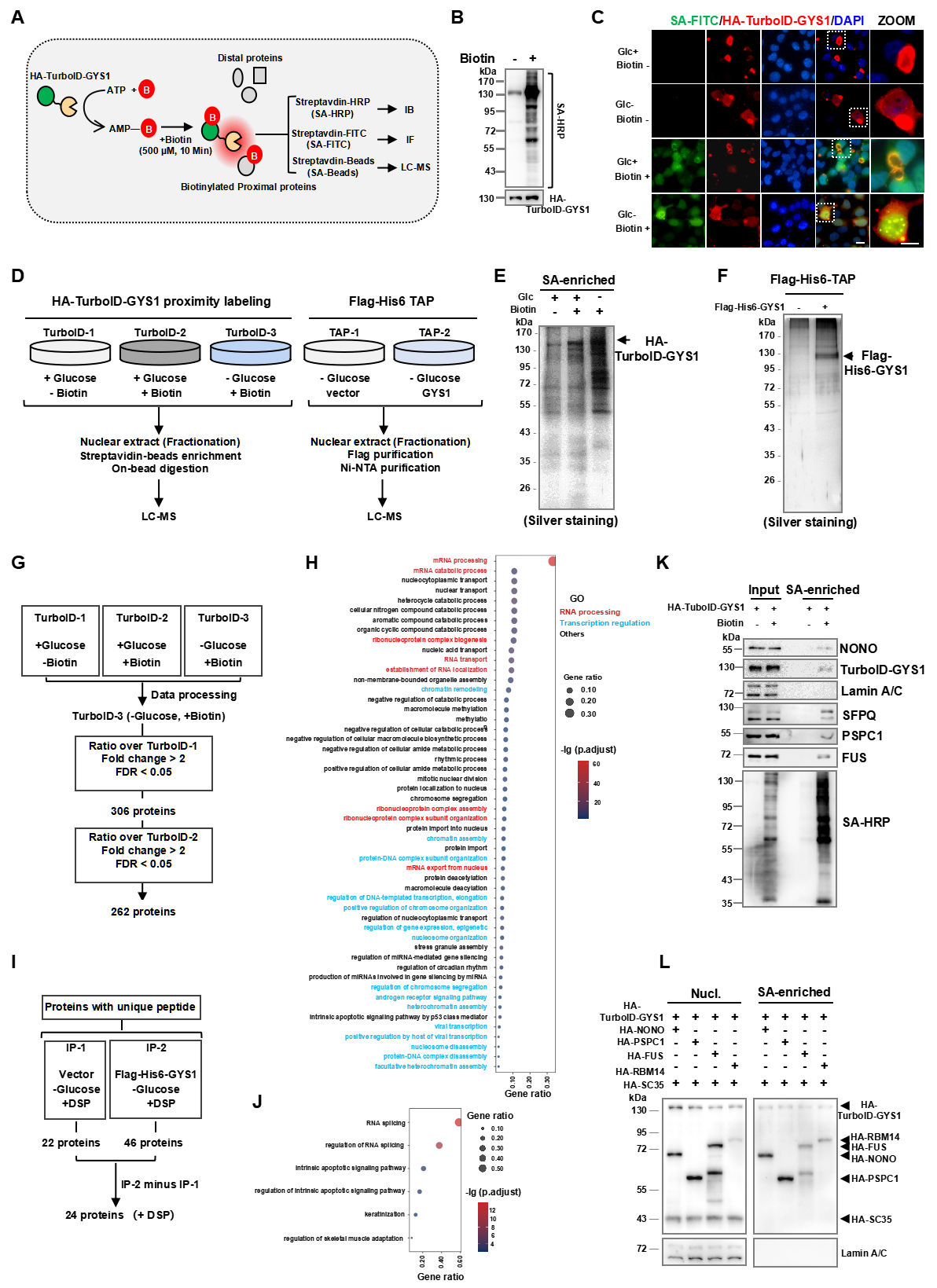

### Fig. S3.tif

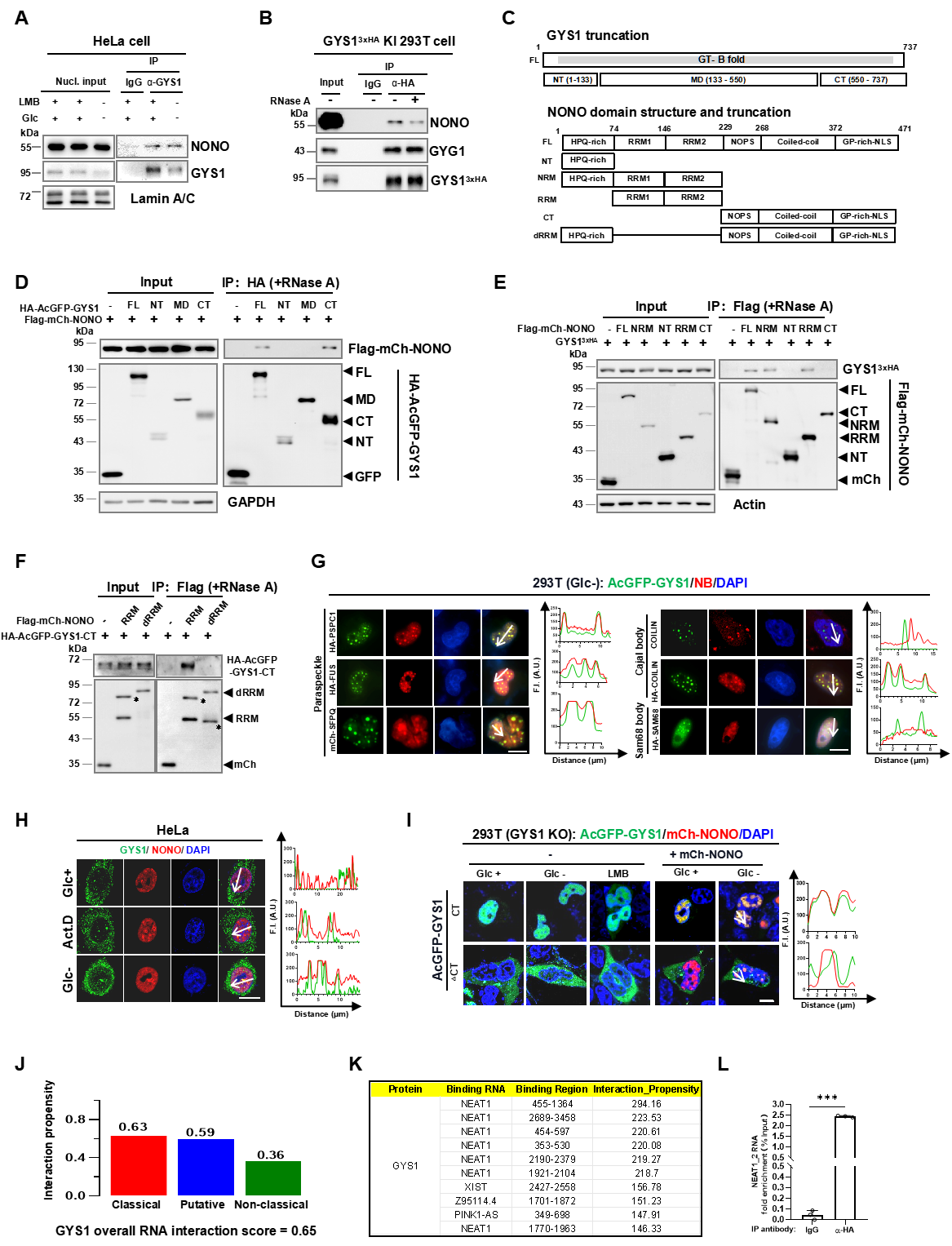

### Fig. S4.tif

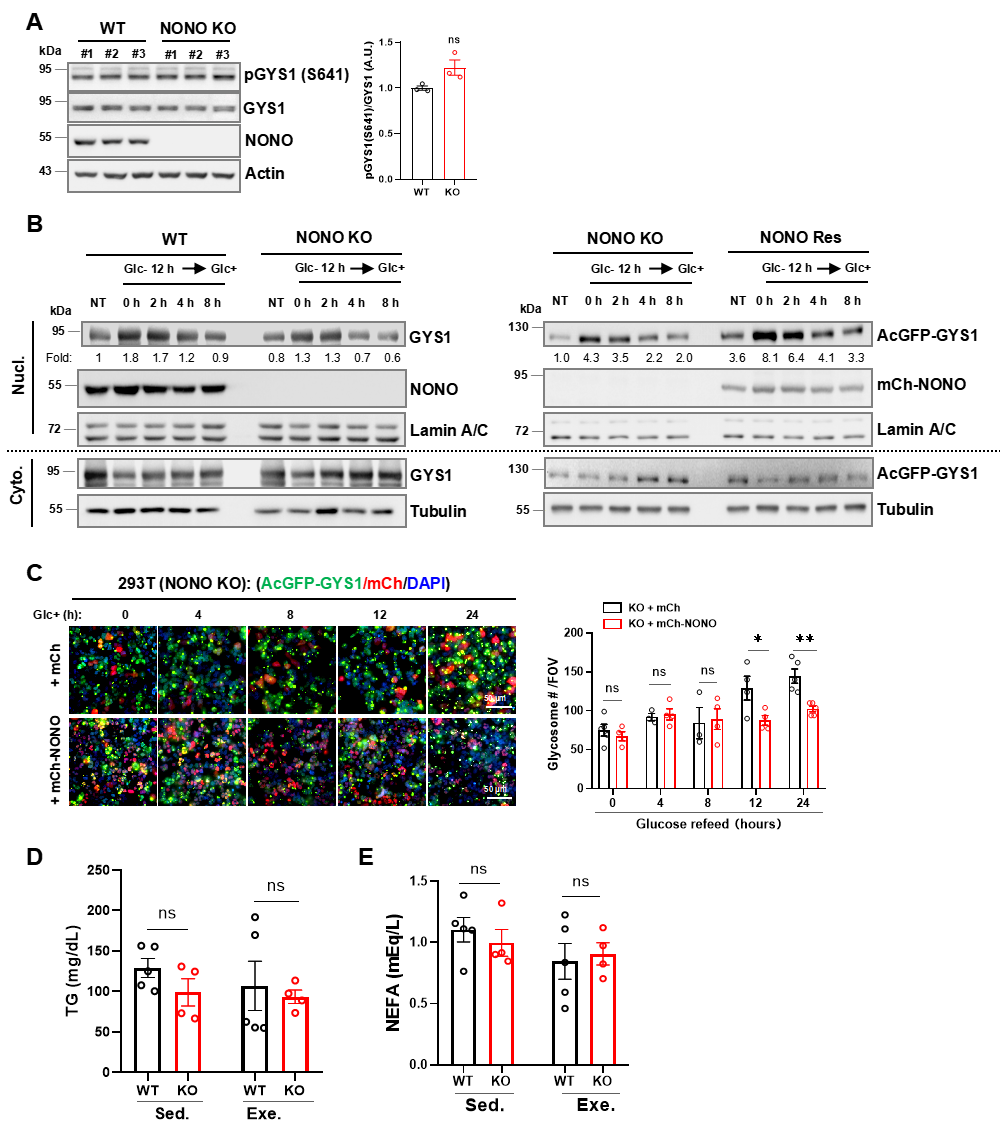

### Fig. S5.tif

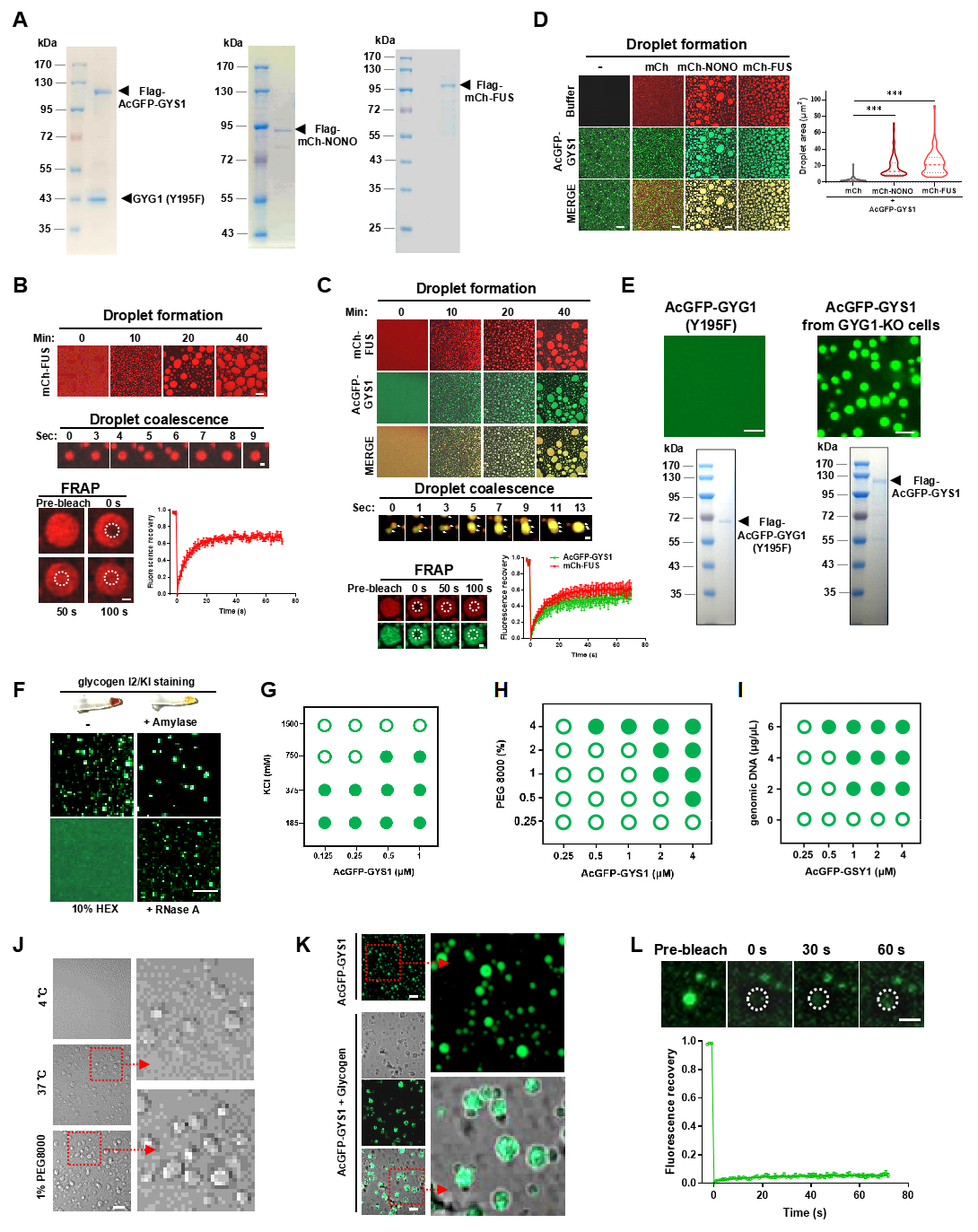

### Fig. S6.tif

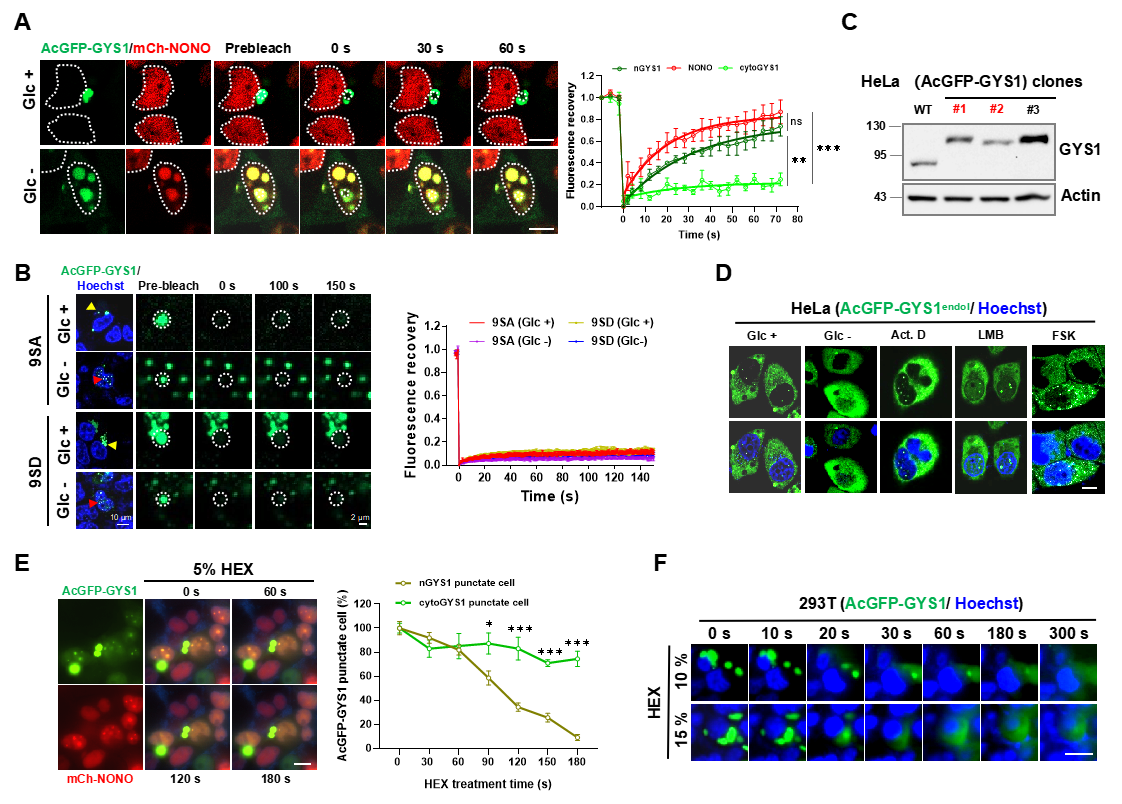

### Fig. S7.tif

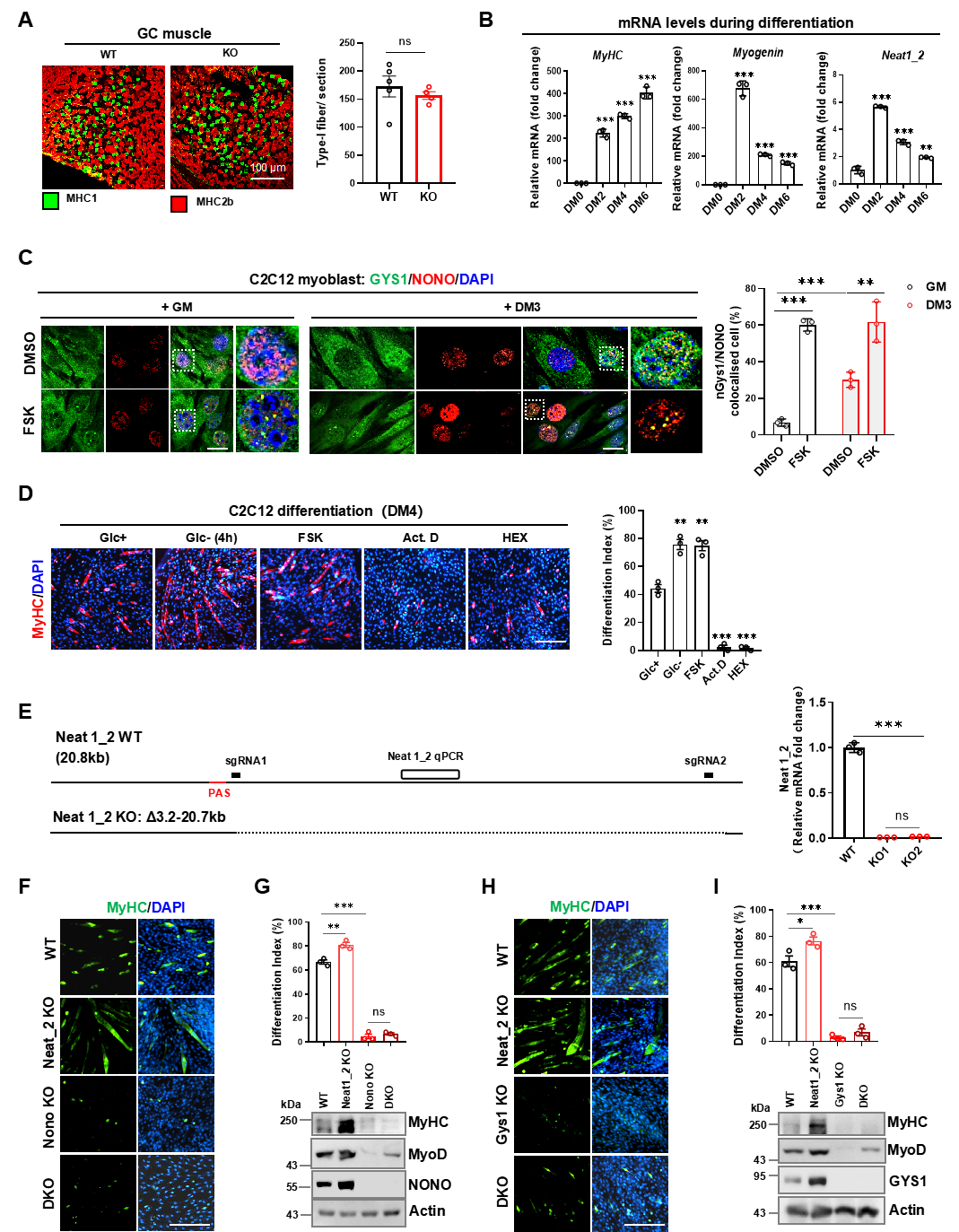

### Fig. S8.tif

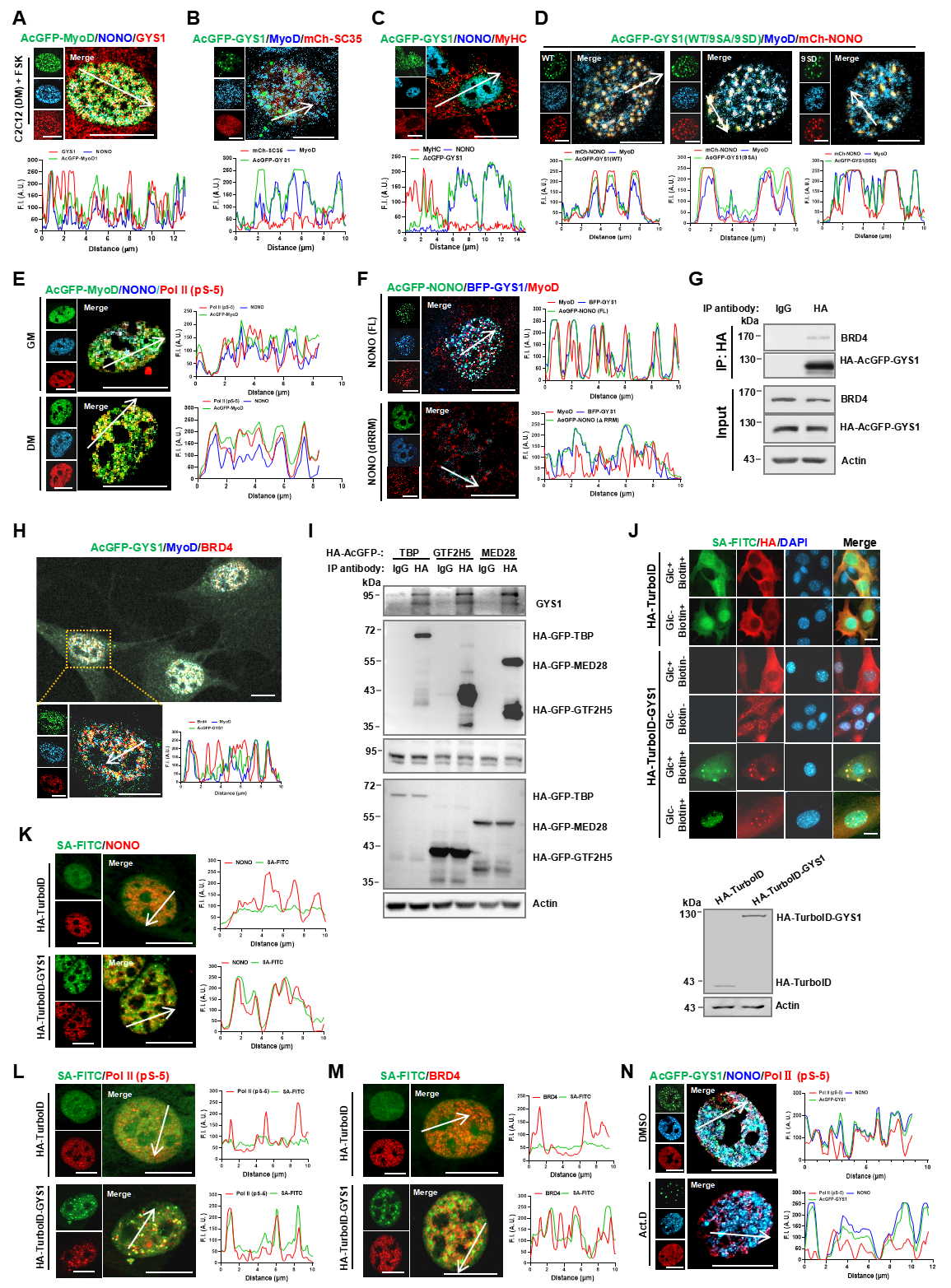

### Fig. S9.tif

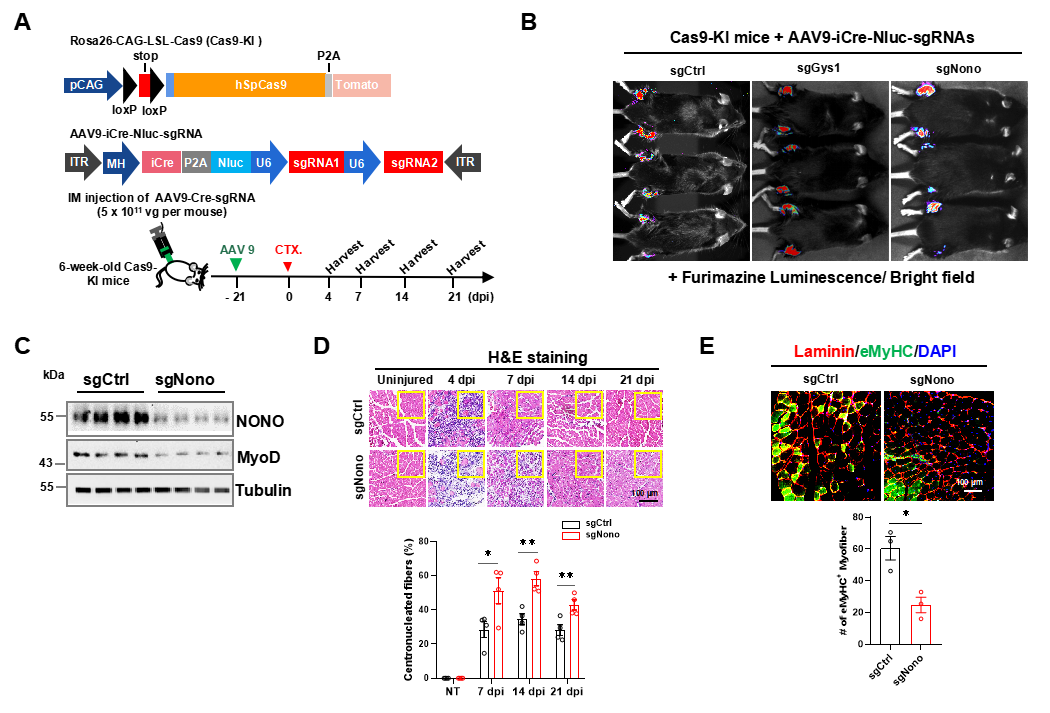
